# Nucleosome patterns in circulating tumor DNA reveal transcriptional regulation of advanced prostate cancer phenotypes

**DOI:** 10.1101/2022.06.21.496879

**Authors:** Navonil De Sarkar, Robert D. Patton, Anna-Lisa Doebley, Brian Hanratty, Adam J. Kreitzman, Jay F. Sarthy, Minjeong Ko, Mohamed Adil, Sandipan Brahma, Michael P. Meers, Derek H. Janssens, Lisa A. Ang, Ilsa Coleman, Arnab Bose, Ruth F. Dumpit, Jared M. Lucas, Talina A. Nunez, Holly M. Nguyen, Heather M. McClure, Colin C. Pritchard, Michael T. Schweizer, Colm Morrissey, Atish D. Choudhury, Sylvan C. Baca, Jacob E. Berchuck, Matthew L. Freedman, Kami Ahmad, Michael C. Haffner, Bruce Montgomery, Eva Corey, Steven Henikoff, Peter S. Nelson, Gavin Ha

**Author notes:** Correspondence: Gavin Ha, Ph.D., Fred Hutchinson Cancer Center 1100 Fairview Ave. N, Seattle, WA 98109; Peter S. Nelson, M.D., Fred Hutchinson Cancer Center 1100 Fairview Ave. N, Seattle, WA 98109. These authors contributed equally (Navonil De Sarkar, Robert D. Patton). Co-senior authors (Peter S. Nelson, Gavin Ha).

## Abstract

Advanced prostate cancers comprise distinct phenotypes, but tumor classification remains clinically challenging. Here, we harnessed circulating tumor DNA (ctDNA) to study tumor phenotypes by ascertaining nucleosome positioning patterns associated with transcription regulation. We sequenced plasma ctDNA whole genomes from patient-derived xenografts representing a spectrum of androgen receptor active (ARPC) and neuroendocrine (NEPC) prostate cancers. Nucleosome patterns associated with transcriptional activity were reflected in ctDNA at regions of genes, promoters, histone modifications, transcription factor binding, and accessible chromatin. We identified the activity of key phenotype-defining transcriptional regulators from ctDNA, including AR, ASCL1, HOXB13, HNF4G, and NR3C1. Using these features, we designed a prediction model which distinguished NEPC from ARPC in patient plasma samples across three clinical cohorts with 97-100% sensitivity and 85-100% specificity. While phenotype classification is typically assessed by immunohistochemistry or transcriptome profiling, we demonstrate that ctDNA provides comparable results with numerous diagnostic advantages for precision oncology.

**STATEMENT OF SIGNIFICANCE:** This study provides key insights into the dynamics of nucleosome positioning and gene regulation associated with cancer phenotypes that can be ascertained from ctDNA. The new methods established for phenotype classification extend the utility of ctDNA beyond assessments of DNA alterations with important implications for molecular diagnostics and precision oncology.

## INTRODUCTION

Metastatic castration-resistant prostate cancer (mCRPC) describes the stage in which the disease has developed resistance to androgen ablation therapies and is lethal (1). Androgen receptor signaling inhibitors (ARSI), designed for the treatment of CRPC, repress androgen receptor (AR) activity and improve survival, but these therapies eventually fail (2, 3). Since the adoption of ARSI as standard-of-care for mCRPC, there has been a prominent increase in the frequency of treatment-resistant tumors with neuroendocrine (NE) differentiation and features of small cell carcinomas (4–7). These aggressive tumors may develop through a resistance mechanism of trans-differentiation from AR-positive adenocarcinoma (ARPC) to NE prostate cancer (NEPC) that lack AR activity (4,7–10). Additional phenotypes can also arise based on expression of AR activity and NE genes, including AR-low prostate cancer (ARLPC) and double-negative prostate cancer (DNPC; AR-null/NE-null) (5,11–13). Distinguishing prostate cancer subtypes has clinical relevance in view of differential responses to therapeutics, but the need for a biopsy to diagnose tumor histology can be challenging: invasive procedures are expensive and accompanied by morbidity, a subset of tumors are not accessible to biopsy, and bone sites pose particular challenges with respect to sample quality (7, 14).

Circulating tumor DNA (ctDNA) released from tumor cells into the blood as cell-free DNA (cfDNA) is a non-invasive “liquid biopsy” solution for accessing tumor molecular information. The analysis of ctDNA to detect mutation and copy-number alterations has served to classify genomic subtypes of CRPC tumors (4,15–21). However, the defining losses of *TP53* and *RB1* in NEPC do not always lead to NE trans-differentiation (7, 22). Rather, ARPC and NEPC tumors are associated with distinct reprogramming of transcriptional regulation (8,9,23). Methylation analysis of cfDNA in mCRPC to profile the epigenome shows promise for distinguishing phenotypes, but requires specialized assays such as bisulfite treatment, enzymatic treatment, or immunoprecipitation (24–27).

The majority of cfDNA represents DNA protected by nucleosomes when released from dying cells into circulation, leading to DNA fragmentation that is reflective of the non-random enzymatic cleavage by nucleases (28, 29). Emerging approaches to analyze cfDNA fragmentation patterns from plasma for studying cancer can be performed directly from standard whole genome sequencing (WGS) (30–34). cfDNA fragments have the characteristic size of 167 bp, consistent with protection by a single core nucleosome octamer and histone linkers, but the size distribution may vary between healthy individuals and cancer patients (35–38). Recent studies have demonstrated that the nucleosome occupancy in cfDNA at the transcription start site (TSS) and transcription factor binding site (TFBS) can be used to infer gene expression and transcription factor (TF) activity from cfDNA (39–41). However, nucleosome positioning and spacing are dynamic in active and repressed gene regulation (42–44). A detailed understanding of the nucleosome organization and positioning patterns associated with transcriptional regulation has not been fully explored in cfDNA.

A major challenge for ctDNA analysis is the low tumor content (tumor fraction) in patient plasma samples. By contrast, plasma from patient-derived xenograft (PDX) models may contain nearly pure human ctDNA after bioinformatic exclusion of mouse DNA reads (36, 38). This provides a resource that is ideal for studying the properties of ctDNA, developing new analytical tools, and validating both genetic and phenotypic features by comparison to matching tumors. In this study, we performed WGS of ctDNA from mouse plasma across 24 CRPC PDX lines with diverse phenotypes. Applying newly developed computational methods, we comprehensively interrogated the nucleosome patterns in ctDNA across genes, regulatory loci, TFBSs, TSSs, and open chromatin sites to reveal transcriptional regulation associated with mCRPC phenotypes. Finally, we designed a probabilistic model to accurately classify treatment-resistant tumors into divergent phenotypes and validated its performance in 159 plasma samples from three mCRPC patient cohorts. Overall, these results highlight that transcriptional regulation of tumor phenotypes can be ascertained from ctDNA and has potential utility for diagnostic applications in cancer precision medicine.

## RESULTS

### Comprehensive resource of matched tumor and liquid biopsies from patient derived xenograft (PDX) models of advanced prostate cancer

We used 26 models from the LuCaP PDX series of advanced prostate cancer with well-defined mCRPC phenotypes (45). The models consisted of 18 classifieds as ARPC, two classified as AR- low and NE-negative prostate cancer (ARLPC), and six classified as NEPC (**Figure 1A, Supplementary Table S1**). For each PDX line, we pooled mouse plasma from seven to ten mice, extracted cfDNA, and performed deep whole genome sequencing (WGS; mean 38.4x coverage, range 21 – 85x) **(Methods, Figure 1A)**. We observed that 25 lines had human ctDNA comprising more than 10% of the sample (mean 52.9%, range 10.6 – 96%) with NEPC samples having significantly higher human fractions (mean 85.1%, range 77.1 – 96%, two-tailed Mann-Whitney U test p = 9.6 x 10^-4^) (**Figure 1B, Supplementary Table S1**). We used bioinformatic subtraction of mouse sequenced reads to obtain nearly pure human ctDNA data **(Methods)**. After subsequent filtering by human ctDNA sequencing coverage, 24 PDX lines remained for further analysis (16 ARPC, 6 NEPC, 2 ARLPC; mean 20.5x, range 3.8 – 50.6x, **Supplementary Table S1**). In the matching tumors, we performed Cleavage Under Targets and Release using Nuclease (CUT&RUN) to profile H3K27ac, H3K4me1, and H3K27me3 histone post-translational modifications (PTMs) (46, 47) (**Supplementary Fig. S1**). We hypothesized that nucleosome organization inferred from ctDNA reflects the transcriptional activity state regulated by histone PTMs (48).

**Figure 1.**
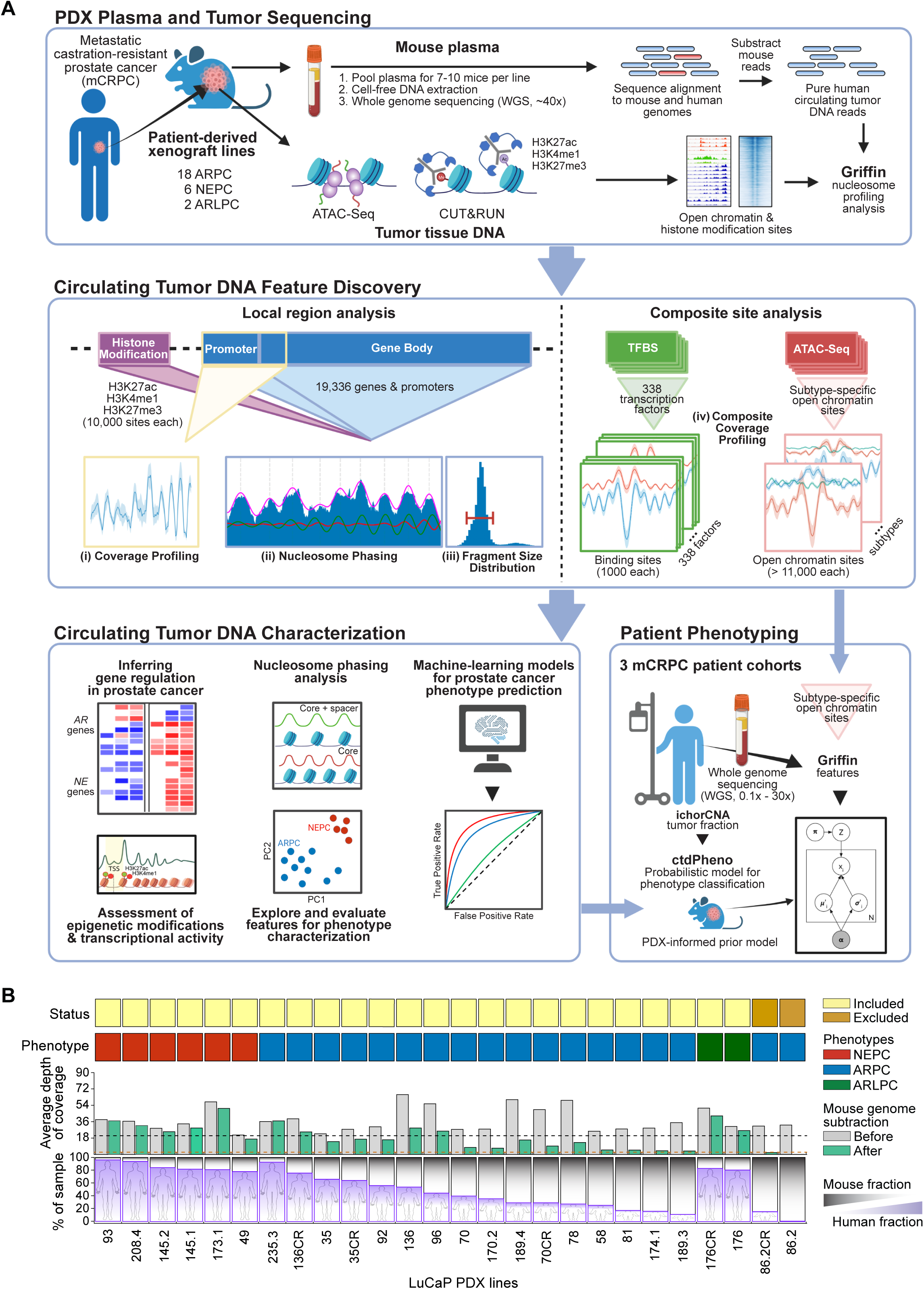
**Characterizing advanced prostate cancer through matched tumor and liquid biopsies from PDX models** (A) (**top**) Both blood and tissue samples were taken from 26 patient-derived xenograft (PDX) mouse models with tumors originating from metastatic castration-resistant prostate cancer (mCRPC) with AR-positive adenocarcinoma (ARPC), neuroendocrine (NEPC), AR-low neuroendocrine-negative (ARLPC) phenotypes. Cell-free DNA (cfDNA) was extracted from pooled plasma collected from 7-10 mice and whole genome sequencing (WGS) was performed. Following bioinformatic mouse read subtraction, pure human circulating tumor DNA (ctDNA) reads remained. From PDX tissue, ATAC-Seq and CUT&RUN (targeting H3K27ac, H3K4me1, and H3K27me3) data were generated. (**middle**) Four distinct ctDNA features were analyzed at five genomic region types using Griffin (49) and nucleosome phasing methods developed in this study (**Methods**). (**bottom**, **left**) Overview of PDX ctDNA features profiled to characterize the mCRPC pathways, transcriptional regulation, and nucleosome positioning. ctDNA features were evaluated for phenotype classification. (**bottom, right**) Phenotype classification using a probabilistic model that accounted for ctDNA tumor content and informed by PDX features was applied to 159 samples in three patient cohorts. (B) PDX phenotypes and mouse plasma sequencing. Inclusion status based on final mean depth after mouse read subtraction (< 3x coverage were excluded; red dotted line). Phenotype status, including 6 NEPC, 18 ARPC (2 excluded), and 2 ARLPC. Average depth of coverage before and after mouse subtraction (mean coverage 20.5x; dotted line). Percentage of the cfDNA sample that contains human ctDNA after mouse read subtraction.

To study transcriptional regulation in mCRPC phenotypes from ctDNA, we interrogated four different features: local promoter coverage, nucleosome positioning, fragment size analysis, and composite TFBSs and open chromatin sites analysis using the Griffin framework (49) (**Figure 1A, Methods**). First, we analyzed three different local regions within ctDNA: all gene promoters and gene bodies and sites of histone PTMs guided by CUT&RUN analysis. For each of the three local regions, we extracted features of nucleosome periodicity using a nucleosome phasing approach and computed the fragment size variability; for promoter regions, we also computed the coverage at the transcription start site (TSS). Next, we analyzed ctDNA at transcription factor binding sites (TFBSs) and open chromatin regions. For each transcription factor (TF), ctDNA coverage at TFBSs were aggregated into composite profiles representing the inferred activity (41, 49). Similarly, features in the composite profiles of subtype-specific open chromatin regions were extracted for analyzing the signatures of chromatin accessibility in ctDNA. Altogether, we assembled a multi-omic sequencing dataset from matching tumor and plasma for a total of 24 PDX lines, making this a unique molecular resource and platform for developing transcriptional regulation signatures of tumor phenotype prediction from ctDNA.

### Characterizing transcriptional activity of AR and ASCL1 in PDX phenotypes through analysis of tumor histone modifications and ctDNA

Prostate cancer phenotypes in mCRPC patients have distinct transcriptional signatures and these are also observed in the LuCaP PDX lines (11). We sought to further characterize the transcriptional activity in different tumor phenotypes by studying epigenetic regulation via histone PTMs. We identified broad peak regions for H3K4me1 (median of 17,643 regions, range 1,894 – 64,934), H3K27ac (median 7,093, range 1610 - 34,047), and H3K27me3 (median 8,737, range 2,024 - 42,495) in the tumors of the 24 PDX lines and an additional nine LuCaP PDX lines where only tumor was available (total of 25 ARPC, 2 ARLPC, and 6 NEPC) (**Methods**, **Supplementary Fig. S1**, **Supplementary Table S2**). Using unsupervised clustering and principal components analysis (PCA), we identified putative active regulatory regions of enhancers and promoters (H3K27ac, H3K4me1) and gene repressive heterochromatic mark (H3K27me3) that were specific to ARPC, ARLPC, and NEPC phenotypes (50) (**Supplementary Fig. S2A**).

AR and ASCL1 are two key differentially expressed TFs with known regulatory roles in ARPC and NEPC phenotypes, respectively (9,51–53). When inspecting AR binding sites in ARPC tumors, we observed increased signals from flanking nucleosomes with H3K27ac PTMs compared to the other phenotypes (area under mean peak profile of 18.46 vs. 15.08 in ARLPC and 10.63 in NEPC) **Figure 2A**, **Supplementary Fig. S2B**, **Methods**). We also observed the strongest signals at the nucleosome depletion region (NDR) in ARPC for H3K27ac (1.54 coverage decrease vs. 0.78 for ARLPC and 0.41 for NEPC). Conversely, in NEPC tumors, we observed stronger signals at nucleosomes with H3K27ac PTMs flanking ASCL1 binding sites (area under mean peak profile 62.65 vs. 29.18 for ARLPC and 10.83 for ARPC), and stronger NDR signals (2.26 coverage decrease vs. 0.19 for ARPC and 0.37 for ARLPC). We observed similar trends for H3K4me1 PTMs in the LuCaP PDX lines (**Supplementary Fig. S2C**).

**Figure 2.**
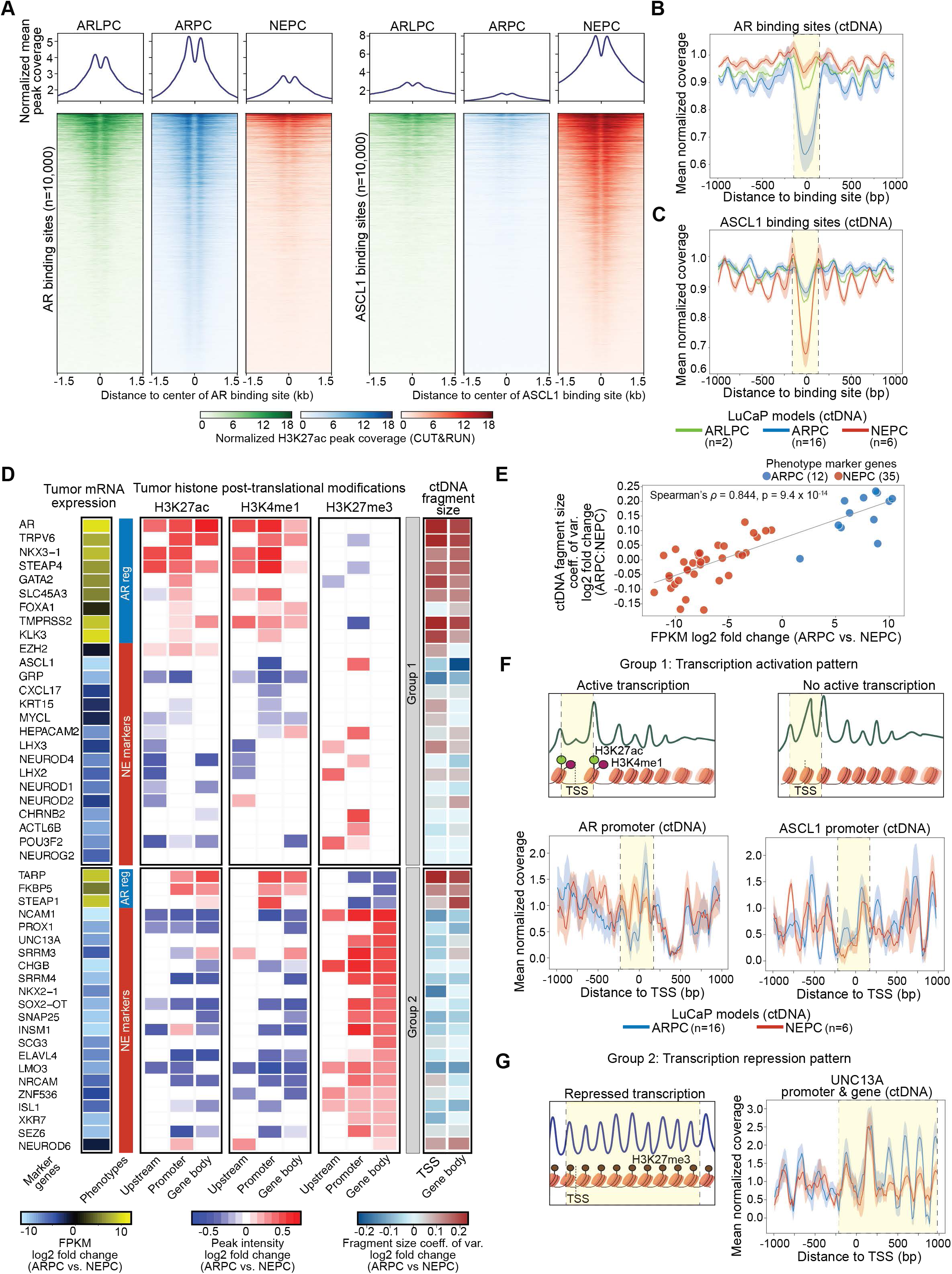
Analysis of tumor histone modifications and ctDNA reveals nucleosome patterns consistent with transcriptional regulation in CRPC phenotype-specific genes. **(A)** H3K27ac peak signals between ARLPC, ARPC, and NEPC PDX tumor phenotypes at 10,000 AR binding sites (left) and at ASCL1 binding sites (right). Binding sites were selected from the GTRD (97) (**Methods**). **(B-C)** Composite coverage profiles at 1000 AR (**B**) and ASCL1 (**C**) binding sites in ctDNA analyzed using Griffin. Coverage profile means (lines) and 95% confidence interval with 1000 bootstraps (shading) are shown. The region ±150 bp is indicated with vertical dotted line and yellow shading. **(D)** Heatmap of log2 fold change in key genes up and down regulated between ARPC and NEPC established through RNA-Seq (**left**) grouped by the type of histone modification which dictates translation levels: Group 1 shows genes where the predominate PTM mark is attributed to H3K27ac or H3K4me1 active marks in the gene promoters or putative distal enhancers, lacking H3K27me3 heterochromatic mark in the gene body; Group 2 features gene body spanning H3K27me3 repression marks. Central columns show differential peak intensity for each of the assayed histone modifications, separated by whether they appear upstream or in the promoter or the body of each gene. On the right the log2 fold change between ARPC and NEPC lines’ fragment size coefficient of variation (CV) is shown for TSS+/1 1KB windows and respective gene bodies. **(E)** Comparison of the log2 fold change (ARPC/NEPC) of mean mRNA expression vs mean coefficient of variation (CV) in the 47 phenotypic lineage marker genes’ promoter regions. **(F)** (**top**) Illustrations of expected ctDNA coverage profiles for Group 1 genes with and without H3K27ac or H3K4me1 modification leading to active and inactive transcription, respectively. (**bottom**) ±1000 bp surrounding the promoter region for AR and ASCL1 in ARPC and NEPC. Shown are coverage profile means (lines) and 95% confidence interval with 1000 bootstraps (shading). Decreased coverage is reflective of increased nucleosome accessibility and thus increased transcription. Dotted line and yellow shading highlight the transcription start site (TSS) at -230 bp and +170 bp. **(G)** Illustration of expected ctDNA coverage profiles for Group 2 genes with repressed transcription caused by H3K27me3 modifications in the gene body. Neuronal gene UNC13A has increased nucleosome phasing in ctDNA of ARPC samples compared to NEPC.

We analyzed the ctDNA composite coverage profiles at TFBSs to evaluate the nucleosome accessibility, whereby lower normalized central (±30 bp window) mean coverage across these sites suggests more nucleosome depletion (**Methods**). For AR TFBSs, we observed the strongest signal for nucleosome depletion in ARPC, as indicated by the lowest mean central coverage (average 0.64, n=16), compared to moderate signals for ARLPC (average 0.88, n=2), and weakest signals for NEPC (average 0.95, n=6) (**Figure 2B**). Conversely, the composite coverage profile at ASCL1 TFBSs showed the strongest nucleosome depletion for NEPC samples (mean central coverage 0.69) compared to ARLPC (0.86) and ARPC (0.88) (**Figure 2C**). These observations were consistent with the differential binding activity by AR and ASCL1 in their respective phenotypes from tumor tissue (**Figure 2A**). Furthermore, the ctDNA coverage patterns of the nucleosome depletion in ctDNA resembled the NDR flanked by nucleosomes with H3K27ac and H3K4me1 peak profiles, which was exemplified when analyzing only nucleosome-sized fragments (140 bp – 200 bp) generated by CUT&RUN (**Figure 2A**, **Supplementary Fig. S2B-C**). Together, these results suggest that the nucleosome depletion in ctDNA at AR and ASCL1 binding sites represents active TF binding and regulatory activity in specific prostate PDX tumor phenotypes.

### Nucleosome patterns at gene promoters inferred from ctDNA are consistent with transcriptional activity for phenotype-specific genes

We selected 47 genes comprising 12 ARPC and 35 NEPC lineage markers established previously (4, 54) and confirmed by differential expression analysis from PDX tumor RNA-Seq data (**Figure 2D**, **Supplementary Table S3**, **Methods**). To assess the activity of these genes from ctDNA, we analyzed the ctDNA fragment size in TSSs (± 1 kb window) and gene bodies, and we found that the differential size variability between phenotypes was positively correlated with relative expression (Spearman’s r = 0.844, p = 9.4 x 10^-14^, **Figure 2E**, **Supplementary Fig. S3A**, **Supplementary Table S2**, **Methods**). Next, we analyzed the relative ctDNA coverage at the TSS (± 1 kb) but did not observe an association between the phenotypes (**Supplementary Fig. S3B)**. However, closer inspection of the ctDNA coverage patterns at the promoters revealed consistent nucleosome organization for transcription activity and repression (39,55–57) (**Figure 2D**). Therefore, we grouped the genes based on differential signals in H3K27me3 histone PTMs, which are associated with repressed transcription or nucleosome compaction (58). For 25 genes (Group 1) without differential H3K27me3 peaks, including AR, FOXA1, KLK3 and ASCL1, we observed nucleosome depletion at the TSS consistent with presence of active PTMs, such as for AR (mean coverage 0.47, n=16) in ARPC and ASCL1 (0.30, n=6) in NEPC samples (**Figure 2F, Supplementary Fig. S4**). By contrast, we observed increased coverage at the TSS of AR (1.08) in NEPC and ASCL1 (0.42) in ARPC, which supports the nucleosome depletion in the absence of PTMs and inactive transcription. For 22 genes (Group 2) with differential H3K27me3 peaks, including STEAP1, CHGB and SRRM4, we observed a relatively more consistent increase in nucleosome occupancy and phasing in the TSS as well as in the gene body for NE-specific genes (**Figure 2G, Supplementary Fig. S5**). The neural signaling genes in this group, such as UNC13A and INSM1, had reduced signals for nucleosome positioning, consistent with the heterogeneous (‘fuzzy’) nucleosome patterns described for actively transcribed genes (43, 59). Interestingly, while UNC13A was repressed in ARPC tumors, it did not have H3K27ac nor H3K4me1 accessible PTM marks in NEPC tumors despite being expressed (**Supplementary Fig. S3B-C**). These results illustrate that ctDNA analysis can reveal patterns that are consistent with transcriptional regulation by histone modifications for key genes that define prostate cancer phenotypes.

### Phasing analysis in ctDNA reveals nucleosome periodicity associated with transcriptional activity between CRPC phenotypes

To systematically quantify inter-nucleosomal spacing and predict nucleosome phasing, we developed TritonNP, a tool utilizing Fourier transforms and band-pass filters on the normalized ctDNA coverage to isolate frequency components with periodicities larger than 146 bp (**Figure 3A**, **Supplementary Fig. S6**, **Methods**). This approach allows for calling phased nucleosome dyad positions to generate an average inter-nucleosome distance from the originating cells, encapsulating potential heterogeneity in nucleosome occupancy and stability. Regions of inactive or repressed transcription are expected to have stably bound nucleosomes, resulting in more periodic (ordered) phasing in the gene body (56,60,61). Conversely, actively transcribed regions may exhibit overall disordered phasing due to transient nucleosome occupancy, resulting in relatively aperiodic patterns with variable degrees of nucleosome depletion. In PDX ctDNA, we observed a larger mean phased-nucleosome distance across 17,946 genes in the ARPC lines compared to the NEPC lines (median 291.1 bp vs. 282.6 bp, p = 0.027; two-tailed Mann-Whitney U test, **Figure 3B**). The phased nucleosome distance was also negatively correlated with the mean cell cycle progression (CCP) score (Spearman’s rho = -0.563, p = 0.006, **Figure 3C**, **Methods**). These results suggest increased nucleosome periodicity in NEPC ctDNA may reflecting the condensed chromatin in hyperchromatic nuclei of NE cells (14), and the phasing analysis may have potential utility for assessing tumor proliferation and aggressiveness (62).

**Figure 3.**
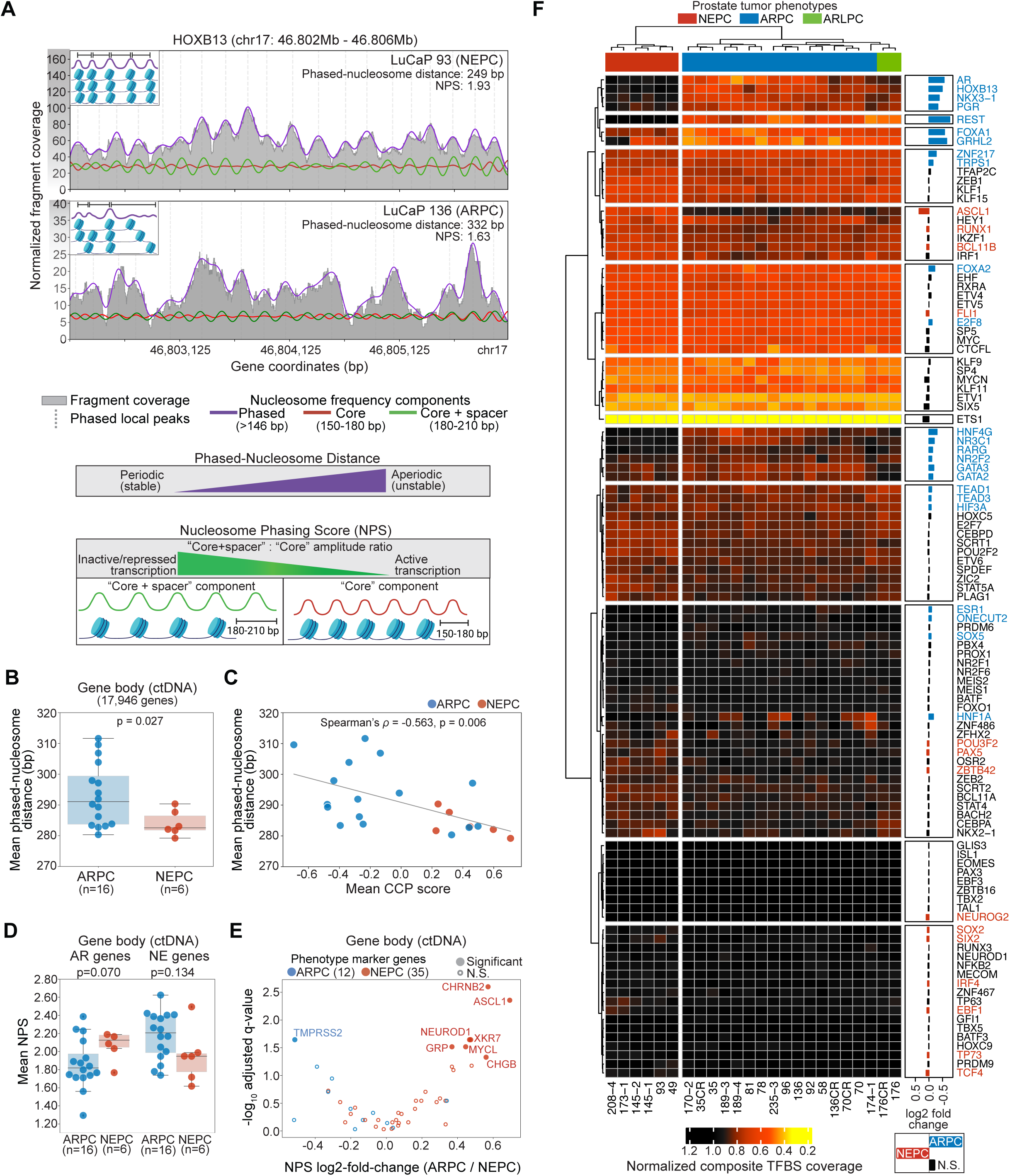
Phasing analysis in ctDNA recapitulates nucleosome stability and trends in transcriptional activity between CRPC phenotypes. **(A)** Illustration of nucleosome phasing analysis using TritonNP for HOXB13, which is expressed in ARPC but not NEPC. Fourier transform and a band-pass filter-based smoothing method was used to phase and identify peaks (grey dotted lines). Frequency components corresponding to > 146 bp (wavelength) are shown in purple. The mean inter-nucleosome distance was computed from all peaks in the gene body: lower values represent more periodic and stable nucleosomes. Nucleosome Phasing Score (NPS) is defined as the ratio of the mean amplitudes between frequency components 180-210 bp (“core + spacer”, green curve) and 150-180 bp (“core”, red curve). **(B)** Boxplot of mean phased-nucleosome distance in 17,946 gene bodies per ctDNA sample for ARPC and NEPC PDX lines. Two-tailed Mann-Whitney U test p-value shown. **(C)** Comparison of the mean phased-nucleosome distance and the mean cell-cycle progression (CCP) score (estimated from RNA-Seq) for 16 ARPC and 6 NEPC PDX lines. **(D)** Boxplot of NPS in gene bodies of 47 phenotype-defining genes (35 NE-regulated and 12 AR-regulated) between ARPC and NEPC lines. Two-tailed Mann-Whitney U test p-values shown. **(E)** Volcano plot of NPS log2-fold-change (ARPC/NEPC) in the 47 phenotype-defining genes. Genes with significantly higher NPS scores (solid-colored dots (two-tailed Mann-Whitney U test, Benjamini-Hochberg adjusted FDR at p < 0.05) and non-significant genes (open circle) are shown. **(F)** Hierarchical clustering of the normalized composite central mean coverage at TFBSs from the Griffin analysis of ctDNA for 107 TFs in LuCaP PDX lines of ARPC (n=16), NEPC (n=6), and ARLPC (n=2) phenotypes. This list of TFs was initially selected as having differential expression between ARPC and NEPC from LuCaP PDX RNA-Seq analysis. Heatmap colors indicate increased accessibility (low values; yellow, orange, red) and decreased accessibility (higher values; black) in ctDNA. TFs with increased accessibility in NEPC samples (log2-fold-change > 0.05, Mann-Whitney U test p < 0.05) are indicated with red text; increased accessibility in ARPC (log2-fold-change < -0.05, p < 0.05) are indicated with blue text.

To model the relationship between nucleosome phasing and transcriptional activity more directly, we further extracted the frequency components corresponding to the inter-nucleosomal distances of the core dyad with spacer (180 – 210 bp) and without (150 – 180 bp). Then, we computed the ratio of the mean frequency amplitudes between these components, called the nucleosome phasing score (NPS), where a higher score corresponded to more ordered nucleosome phasing and repressed transcription. As an example, HOXB13, which is transcriptionally inactive in NEPC had higher overall GC-corrected coverage (mean 56.85 depth) and a phased nucleosome distance of 249 bp with a 1.93 NPS in the LuCaP 93 NEPC PDX (**Figure 3A**). By contrast, HOXB13 is actively transcribed in ARPC and had lower coverage (mean 13.54 depth) and a more disordered phased-nucleosome distance of 332 bp with a 1.63 NPS in the LuCaP 136 ARPC PDX. When assessing the 47-prostate cancer phenotype marker genes, we observed that the mean NPS for the 35 NE genes was lower in NEPC lines compared to ARPC (median NPS 1.95 vs. 2.21, p = 0.134; two-tailed Mann-Whitney U test, **Figure 3D**); although this was not statistically significant, it was consistent with their active transcription. Conversely, the mean NPS for the 12 AR-regulated genes was lower in ARPC lines compared to NEPC (median NPS 1.82 vs 2.13, p = 0.070; two-tailed Mann-Whitney U test). In particular, 26 (74%) of the NE genes had lower NPS in NEPC compared to ARPC (log2 fold-change [ARPC:NEPC] > 0), including seven genes (ASCL1, CHGB, CHRNB2, GRP, MYCL, XKR7, NEUROD1) that were statistically significant (p < 0.05); ten (83%) of the AR-regulated genes had lower NPS in ARPC (log2 fold-change < 0), with TMPRSS2 being statistically significant (**Figure 3E**, **Supplementary Table S3**). These results illustrate that the NPS captured signals distinguishing key lineage-specific gene markers.

### Inferred TF activity from analysis of nucleosome accessibility at TFBSs in ctDNA confirms key regulators of tumor phenotypes

To characterize the lineage-defining TFs in prostate tumor phenotypes, we considered nucleosome accessibility at TFBSs in PDX ctDNA. We identified 107 TFs based on the intersection of 338 TFs analyzed using Griffin and 404 differentially expressed TFs between ARPC and NEPC PDX tumors (**Supplementary Fig. S7, Supplementary Table S3, Methods**). Of these TFs, 38 had significantly different accessibility in ctDNA between ARPC and NEPC phenotypes (two tailed Mann-Whitney U test, Benjamini-Hochberg adjusted p < 0.05, **Supplementary Table S3**). Through unsupervised hierarchical clustering of composite TFBS central coverage values for the 107 TFs, we observed distinct groups of TFs in PDX ctDNA (**Figure 3F**). REST had the largest difference in accessibility as supported by a decrease in coverage within ARPC models compared to NEPC (log2 fold-change -0.77, adjusted p = 5.7 x 10^-4^, **Supplementary Fig. S8A**, **Supplementary Table S3**). FOXA1, and GRHL2 were significantly more accessible in ARPC (and ARLPC) samples compared to NEPC (log2 fold-change < -0.57, adjusted p < 1.3 x 10^-3^). AR, HOXB13, and NKX3-1 had higher accessibility in ARPC compared to NEPC (log2 fold-change < -0.37, adjusted p < 1.3 x 10^-3^), but with only moderate accessibility in ARLPC, as expected. Interestingly, progesterone receptor (PGR) also had high accessibility in ARPC (log2 fold-change -0.33, adjusted p = 2.6 x 10^-3^, **Supplementary Fig. S8A)**. We also observed a group of ARPC-regulated genes that followed a similar trend, including the glucocorticoid receptor (NR3C1) and other nuclear hormone receptors (NR2F2, RARG), pioneer factors GATA2 and GATA3, and nuclear factors HNF4G and HNF1A (log2 fold-change < -0.10, adjusted p < 0.027, **Supplementary Fig. S8A**).

For factors that had higher accessibility in NEPC models compared to ARPC and ARLPC, ASCL1 had the largest TFBS coverage difference (log2 fold-change 0.36, adjusted p = 5.7 x 10^-4^, **Figure 2C**, **Figure 3F)**. Other TFs, including RUNX1, BCL11B, POU3F2, NEUROG2, and SOX2 also had higher activity in NEPC (log2 fold-change > 0.06, adjusted p < 0.048, **Supplementary Fig. S8B**), although the difference was modest. HEY1, IRF1, and IKZF1 had a similar trend consistent with increased accessibility in NEPC samples but were not significantly different from ARPC (adjusted p > 0.10). While NKX2-1 and CEBPA had increased accessibility in NEPC compared to ARPC (although not significant with adjusted p = 0.47 and 0.36 respectively), these factors were also modestly active in ARLPC (**Supplementary Fig. S8B, Supplementary Table S3**). Other notable factors such as MYC and ETS transcription family genes (ETV4, ETV5, ETS1, ETV1) had high accessibility across all phenotypes, while NEUROD1, RUNX3, and TP63 were inaccessible in nearly all samples. Overall, we identified the accessibility of known prostate cancer regulators, including ASCL1, NR3C1, HNF4G, HNF1A, and SOX2 (63–65), that have not been shown before from ctDNA analysis in these tumor phenotypes.

### Phenotype-specific open chromatin regions in PDX tumor tissue are reflected in ctDNA profiles of nucleosome accessibility

Nucleosome profiling from cfDNA sequencing analysis has shown agreement with overall chromatin accessibility in tumor tissue (37,41,66); however, its application for distinguishing tumor phenotypes has been limited. We investigated the use of ATAC-Seq data from tumor tissue for 10 LuCaP PDX lines (5 ARPC and 5 NEPC) to inform phenotype differences in chromatin accessibility (9). We defined an initial set of 28,765 ARPC and 21,963 NEPC differential consensus open chromatin regions which we further restricted to those that overlapped TFBSs for 338 TFs, resulting in 15,881 ARPC and 11,694 NEPC sites (**Methods**, **Figure 4A**). For ARPC- specific open chromatin sites, we observed decreased overall composite site coverage (+/- 1 kb window) and central coverage (+/- 30 bp) in the ctDNA for ARPC PDX lines (mean central coverage 0.75, n=16) compared to NEPC lines (mean 0.96, n=6) and cfDNA from healthy human donors (mean 0.97, n=14) (**Figure 4B**, **Supplementary Table S3**). Conversely, for NEPC-specific open chromatin sites, coverage was decreased in ctDNA for NEPC lines (mean 0.89) compared to ARPC lines (mean 1.01) and healthy donors cfDNA (mean 1.00) (**Figure 4C**, **Supplementary Table S3**). These results confirmed that tumor tissue chromatin accessibility can be corroborated in ctDNA and that ARPC and NEPC phenotypes have distinct ctDNA composite site coverage profiles.

**Figure 4.**
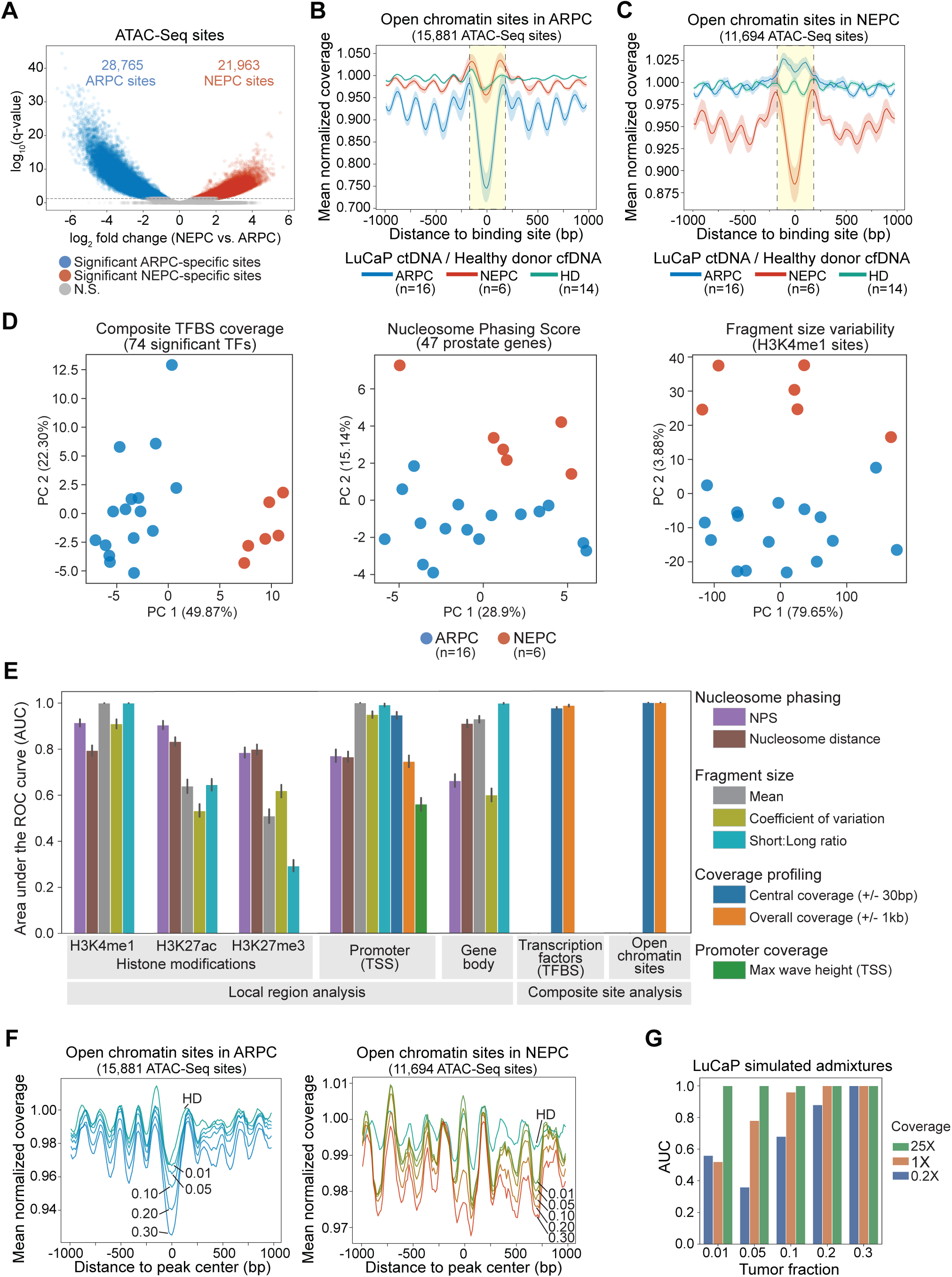
**Comprehensive evaluation of ctDNA features throughout the genome for CRPC phenotype classification in PDX models** (A) Volcano plot of log2-fold change of ATAC-Seq peak intensity between 5 ARPC and 5 NEPC lines; the dotted line demarcates sites by q-value < 0.05. **(B-C)** Composite coverage profiles at open chromatin sites specific to ARPC (**B**) and NEPC (C) PDX tumors analyzed by Griffin. Sites from (A) were filtered for overlap with known TFBSs in 338 factors from GTRD (97). Coverage profile means (lines) and 95% confidence interval with 1000 bootstraps (shading) are shown. The region ±150 bp is indicated with vertical dotted line and yellow shading. (D) PCAs of ctDNA features demonstrates grouping between ARPC and NEPC phenotypes: (**left**) Composite central coverage of TFBSs significant for 74 TFs with differential accessibility out of 338 factors between ARPC and NEPC (**Supplementary Table S4**). (**center**) NPS in the gene bodies of the 47 phenotype defining genes. (**right**) Fragment size variability (coefficient of variation) at H3K4me1 histone modification sites (n=9,750). (E) Performance of classifying ARPC vs NEPC PDX from ctDNA using supervised machine learning (XGBoost) in various region types (all genes, TFBSs, and open regions, **Methods**). Area under the receiver operating characteristic curve (AUC) with 95% confidence interval (100 repeats of stratified cross validation) is shown for performance of all feature types. (F) Example composite coverage profiles at open chromatin sites specific to ARPC (left) and NEPC (right) identified in **B**-**C**. Simulated admixtures generated using ARPC mixed with healthy donor (HD) (left) and NEPC mixed with HD (right) are shown for varying tumor fractions. (G) Performance for classification on admixtures samples using ctdPheno. Five ctDNA admixtures were generated for each phenotype from PDX lines, each at various sequencing coverages and tumor fractions. In total, 125 admixtures were evaluated. The mean AUC across the 5 admixtures is shown for each configuration.

### Comprehensive evaluation of ctDNA features across genomic contexts for CRPC phenotype classification

To assess the utility of ctDNA nucleosome profiling for informing prostate cancer phenotype classification, we systematically evaluated four groups of global genome-wide ctDNA features: phasing, fragment sizes, local coverage profiling, and composite site coverage profiling (**Figure 1A**). From principal components analysis (PCA), we observed distinct feature signals between ARPC and NEPC phenotypes for composite TFBS coverage of TFs, NPS of 47 phenotype marker genes, and fragment size variability at global sites of PTMs (**Figure 4D**, **Supplementary Fig. S9A, Supplementary Table S4**, **Methods**). In addition to these features, we also included similar approaches previously reported, including short-long fragment ratio and local coverage patterns at the TSS (max wave height between -120bp to 195bp) (30, 40) (**Methods**).

We quantitatively evaluated all combinations of coverage, phasing, and fragment size features for different genomic contexts to investigate their potential to classify ARPC and NEPC phenotypes. For each feature set, we conducted 100 iterations of stratified cross-validation using a supervised machine learning classifier (XGBoost) on ctDNA samples from the 16 ARPC and 6 NEPC models and computed the area under the receiver operating characteristic curve (AUC) (**Methods**). First, we evaluated an established set of 10 genes associated with AR activity (5, 12). We observed that the phased nucleosome distance at H3K27ac sites and the central coverage at TSSs had moderate predictive performance (AUC 0.88) (**Supplementary Fig. S9B**, **Supplementary Table S4**). For the set of 47 phenotype markers, the NPS of gene bodies was most predictive (AUC 0.98) (**Supplementary Fig. S9C Supplementary Table S4)**. When considering all PTM sites, promoters, genes, TFs, and open chromatin regions, the best performing features included mean fragment size at H3K4me1 sites (n=9,750, AUC 1.0) and promoter TSSs (n=17,946, AUC 1.0), and both open chromatin composite site features (AUC 1.0) (**Figure 4E**, **Supplementary Table S4**).

### Accurate classification of ARPC and NEPC phenotypes from patient plasma using a probabilistic model informed by PDX ctDNA analysis

An important consideration and challenge in analyzing plasma from patients is the presence of cfDNA released by hematopoietic cells, which leads to a lower ctDNA fraction (i.e., tumor fraction). Furthermore, the small patient cohorts with available tumor phenotype information make supervised machine learning approaches suboptimal. Therefore, we developed ctdPheno, a probabilistic model to estimate the proportion of ARPC and NEPC from an individual plasma sample, accounting for the tumor fraction (**Methods**). We focused on the phenotype-specific open chromatin composite site features and used the PDX plasma ctDNA signals (**Figure 4B-C**, **Supplementary Table S3**) to inform the model. The model produces a normalized prediction score that represents the estimated signature of ARPC (lower values) and NEPC (higher values). We applied this method to benchmarking datasets generated by simulating varying tumor fractions and sequencing coverages using five ARPC and NEPC PDX ctDNA samples each (**Figure 4F**, **Methods**). We achieved a 1.0 AUC at 25X coverage down to 0.01 tumor fraction, 1.0 AUC at 1X down to 0.2 tumor fraction, and 1.0 AUC at 0.2x coverage at 0.3 tumor fraction, suggesting a possible upper-bound performance for classifying samples with lower tumor fraction in plasma (**Figure 4G**, **Supplementary Table S4**).

To test the classification performance of the model on patient samples, we analyzed a published dataset of ultra-low-pass whole genome sequencing (ULP-WGS) of plasma cfDNA (mean coverage 0.52X, range 0.28-0.92X) from 101 mCRPC patients comprising 80 adenocarcinoma (ARPC) and 21 NEPC samples (DFCI cohort I) (25). Using the model, which was unsupervised and used parameters informed only by the PDX analysis, we achieved an overall AUC of 0.96 (**Figure 5A**, **Supplementary Table S5**). When considering samples with high (≥ 0.1) and low (< 0.1) tumor fraction, the model had an 0.97 AUC and 0.76 AUC, respectively (**Supplementary Fig. S10A**). We identified an optimal overall performance at 97.5% specificity (ARPC) and 90.4% sensitivity (NEPC) which corresponded to the prediction score of 0.3314 (**Figure 5A**). These results were concordant (92.1%) with phenotype classification by cfDNA methylation on the same plasma samples (**Supplementary Fig. S10B**, **Supplementary Table S5**). In another published dataset of 11 mCRPC samples from 6 patients who had high PSA, treatment with ARSI, or both (DFCI cohort II) (67, 68), the model correctly classified patients as ARPC in 11 (100%) WGS (∼20x) and 8 (73%) ULP-WGS (∼0.1x) samples when using the optimal score cutoff (**Figure 5B**, **Supplementary Table S5**).

**Figure 5.**
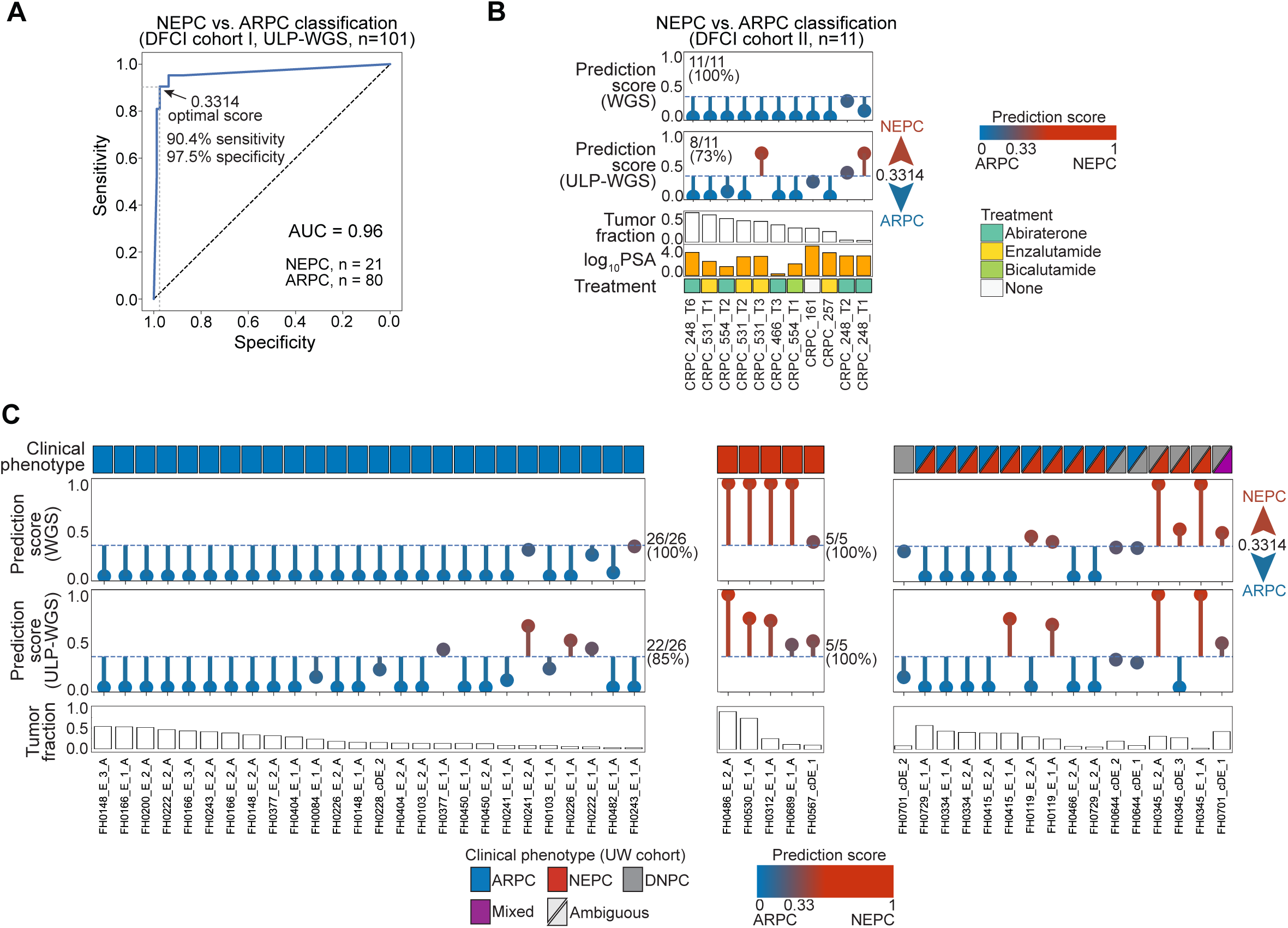
**Accurate classification of NEPC phenotypes from plasma in three patient cohorts using a probabilistic model (ctdPheno) informed by PDX ctDNA features** (A) Receiver operating characteristic (ROC) curve for 101 mCRPC patients (DFCI cohort I) with ultra-low-pass WGS (ULP-WGS) data. The optimal performance of 90.4% sensitivity (for predicting NEPC) and 97.5% specificity (for predicting ARPC) corresponding to a prediction score cutoff of 0.3314 is indicated with horizontal and vertical dotted lines, respectively. (B) Prediction scores for 11 plasma samples from seven patients (DFCI cohort II) with both WGS and ULP-WGS data. The 0.3314 score cutoff threshold (dotted line) was used for classifying NEPC and ARPC. Tumor fractions were estimated by ichorCNA from WGS data. Patients were treated for adenocarcinoma (ARPC) or had high PSA values. (C) Prediction scores for 47 plasma samples with clinical phenotypes comprising 26 ARPC (blue), 5 NEPC (red), and 16 mixed or ambiguous phenotypes (purple, triangles), including double-negative prostate cancer (DNPC; grey). Scores are shown for WGS and ULP-WGS (0.1X) for the same ctDNA sample. The cutoff threshold of 0.3314 (dotted line) was used for classifying NEPC and ARPC. Tumor fractions were estimated by ichorCNA on the WGS data.

Next, we analyzed 61 clinical plasma samples from 30 CRPC patients with ARPC, NEPC, and mixed phenotypes that are representative of typical clinical histories (**Supplementary Table S5**). We performed ULP-WGS of cfDNA and selected 47 samples from 30 patients (26 ARPC, 5 NEPC, and 16 mixed phenotypes) based on tumor fraction and AR copy number status and performed deeper WGS (mean 22.13X coverage, range 15.15X – 31.79X) (**Supplementary Table S5**, **Methods**). For the 26 samples with ARPC clinical phenotype, we predicted all to be predominantly ARPC using the score cutoff of 0.3314 (**Figure 5C**). For NEPC clinical phenotype, all five were predicted to be NEPC with scores above the cutoff. We also noted a negative association between the patient ctDNA coverage at open chromatin sites and the tumor fraction for both ARPC (Spearman’s r = -0.93) and NEPC predictions (Spearman’s r = -1.00), suggesting that the observed ctDNA signals were likely tumor-specific (**Supplementary Fig. S10C**). From ULP-WGS data, we correctly predicted 22 (84%) samples with ARPC clinical phenotype and all five (100%) samples with NEPC clinical phenotype (**Figure 5C**). The remaining 16 samples had clinical histories or tumor histologies that reflected mixed phenotypes such as a tumor with AR-positive adenocarcinoma intermixed with NEPC (**Figure 5C**, **Supplementary Table S5, Supplementary Fig. S11**). For 12 samples that included presence of ARPC in the mixed clinical phenotype, 10 (83%) were classified as ARPC at the optimal score cutoff. For all three samples that had presence of NEPC but no ARPC in the clinical phenotype, the model classified them as NEPC (**Supplementary Fig. S12**). Overall, we achieved an accuracy of 100% for WGS (87% for ULP-WGS) data for samples with unambiguous clinical phenotypes. However, the variable predictions for mixed or ambiguous phenotypes underscore the complexities associated with classification in patients with advanced prostate cancer where tumor heterogeneity can be observed.

## DISCUSSION

To our knowledge, we present the largest sequencing study to date of human ctDNA from mouse plasma of PDX models. The sequencing of mouse plasma provided a unique opportunity to comprehensively interrogate the epigenetic nucleosome patterns in ctDNA from well-characterized tumor models. We developed and applied computational methodologies to construct a multitude of ctDNA features, each of which were associated with the transcriptional regulation in the LuCaP PDX models across CRPC tumor phenotypes. Using features learned from the PDX ctDNA, we developed a probabilistic model to accurately classify ARPC and NEPC phenotypes from patient plasma in three clinical cohorts.

The use of PDX mouse plasma overcomes the challenge of low ctDNA content or incomplete knowledge of the tumor when studying patient samples and can expedite development of cfDNA diagnostics, basic cancer research, and clinical translation. Furthermore, the LuCaP ctDNA sequencing data complements the maturing characterization of CRPC tumor phenotypes from tissue. In addition to supporting molecular studies of CRPC, the ctDNA data and our approaches expand on the potential utility of PDX models for translational research. While these data were focused on ARPC and NEPC phenotypes, this study may serve as a framework for the use of PDX plasma from additional CRPC phenotypes and other cancers models.

The analysis of the LuCaP PDX ctDNA sequencing data confirmed the activity of key regulators between ARPC and NEPC phenotypes, including a set of 47 established differentially expressed gene markers. While gene expression inference from ctDNA has been shown in proof-of-concept studies (34, 40), the PDX ctDNA allowed for a detailed dissection of nucleosome organization associated with transcriptional activity of individual genes that define the tumor phenotypes. Previous analytical approaches have profiled nucleosome occupancy from cfDNA (37, 66). However, our assessment of nucleosome stability by means of the Nucleosome Phasing Score is the first to capture the highly variable spacing and position of the nucleosome arrays associated with transcription and tumor aggressiveness (42,62,69).

In addition to the existing molecular profiling available for these models, we now provide characterization of histone PTMs in LuCaP PDX tumors using CUT&RUN. At regions with these PTMs on histone tails, we observed expected nucleosome patterns inferred in ctDNA that were consistent with active or repressed gene transcription. To our knowledge, this is the first time that ctDNA analysis has been performed in the context of histone PTMs and will provide a blueprint to develop new approaches for studying additional epigenetic alterations using PDX plasma.

While the regulation of key factors such as AR, HOXB13, NKX-3.1, FOXA1, and REST has been shown from ctDNA in CRPC (41), we report the differential activity of other key factors in CRPC for the first time from ctDNA analysis. This included the glucocorticoid receptor (NR3C1), nuclear factors HNF4G and HNF1A, and pioneering factors GATA2 and GATA3, all of which are associated with prostate adenocarcinoma (ARPC) (63,65,70). ASCL1 is a pioneer TF with roles in neuronal differentiation and was recently described to be active during NE trans-differentiation and in NEPC (9, 53). To our knowledge, this study is the first to demonstrate ASCL1 binding site accessibility and provide a detailed characterization of its transcriptional activity in NEPC from plasma ctDNA.

We show an expansive analysis of TFBSs for 338 factors in each plasma sample without the need for chromatin immunoprecipitation or other epigenetic assays. However, we did not find a significant difference in accessibility for 69 out of the 107 TFs in ctDNA, which may be consistent with TF activity not necessarily being correlated with its own expression levels (71). On the other hand, the accessibility of TFBSs may not necessarily indicate true TF activity, such as binding of multiple factors to the same locus. Moreover, our analysis was based on TFBSs obtained from public databases; however, prostate phenotype-specific TF cistromes can better guide this approach.

We applied state-of-the-art computational approaches built on existing and new concepts of ctDNA data analysis to extract tumor-specific features, including the representation of nucleosome phasing, periodicity, and spacing associated with transcriptional activity. Other approaches have also considered regions, such as TSSs, TFBSs, and DNase hypersensitivity sites (33,37,40,41); however, after a systematic evaluation, we found that ctDNA features in open chromatin sites derived from ATAC-Seq of PDX tissue (9) provided the highest performance for distinguishing CRPC phenotypes. We presented ctdPheno which is an unsupervised probabilistic model that estimates the proportion of ARPC and NEPC in patient plasma using a statistical framework informed by idealized parameters from the LuCaP PDX ctDNA analysis. This model does not require training on patient samples but does require tumor fraction estimates (ichorCNA (72)) and a prediction score cutoff determined from DFCI cohort I. Another current limitation is the prediction of only ARPC and NEPC phenotypes; however, the framework can be extended to model multiple phenotype classes, provided the informative parameters for these additional states can be learned. Insights from additional datasets such as single-cell nucleosome and accessibility profiling (73, 74) of PDX tumors and clinical samples may improve the resolution for ctDNA analysis.

Applying the prediction model to patient datasets with definitive clinical phenotypes yielded high performance despite using low depth of coverage sequencing. In particular, our performance for the DFCI cohort I was also consistent with the reported phenotype classification results using ctDNA methylation in the same patients (25). Similarly, in the UW cohort, samples with well-defined clinical phenotypes had perfect concordance from deep WGS data. However, samples with mixed or ambiguous clinical phenotypes limited our ability to definitively assess the performance of the model because a subset of cases had complex clinical and histopathological features. Tumor heterogeneity and co-existence of different molecular phenotypes are common in mCRPC where treatment-induced phenotypic plasticity may vary within and between tumors in an individual patient. Larger studies with comprehensive assessment of the tumor histologies will be needed for developing future extensions of the model to predict mixed phenotypes from ctDNA.

In summary, this study illustrates for the first time that analysis of ctDNA from PDX mouse plasma at scale can facilitate a more detailed investigation of tumor regulation. These results, together with the suite of computational methods presented here, highlight the utility of ctDNA for surveying transcriptional regulation of tumor phenotypes and its potential diagnostic applications in cancer precision medicine.

## ACKNOWLEDGEMENTS

We thank the many patients and their families for their altruistic contributions to this study. We thank the Fred Hutchinson Cancer Research Center Genomics Shared Resources Core members, the Institute for Prostate Cancer Research clinicians and staff that support the University of Washington rapid autopsy program and the PDX program. We thank Patricia Galipeau and members of the Ha and Nelson Laboratories for critically reading this manuscript. We also thank Srinivas Ramachandran for critical discussion and helpful advice in the preliminary phases of this work.

## AUTHOR CONTRIBUTIONS

Conceptualization: N.D.S., R.D.P., G.H., P.S.N.

Methodology: N.D.S., R.D.P., A-L.D., P.S.N., G.H.

Software: R.D.P., A-L.D., N.D.S., B.H., A.J.K., G.H.

Formal Analysis: R.D.P., N.D.S., A-L.D., B.H., A.J.K., M.K., M.A., I.C., G.H.

Investigation: N.D.S., R.D.P., A-L.D., B.H., J.F.S., J.M.L., A.B., G.H.

Resources: N.D.S., J.S., S.B., M.P.M., D.H.J., L.A.A., R.F.D., T.A.N., H.M.M., S.C.B., J.E.B., M.L.F., C.M., H.M.N., E.C., S.H., P.S.N., G.H.

Data Curation: N.D.S., R.D.P., A-L.D., M.P.M., S.B., D.H.J., E.C., C.M., A.D.C., M.C.H., P.S.N., G.H.

Writing – Original Draft: R.D.P, N.D.S, P.S.N., G.H.

Writing – Review & Editing: N.D.S, R.D.P., A-L.D., C.C.P., C.M., A.D.C., M.T.S., B.M., M.C.H., E.C., K.A., S.H., P.S.N., G.H.

Visualization: R.D.P., N.D.S., B.H., M.K., M.C.H., P.S.N., G.H.

Supervision: P.S.N., G.H. Funding Acquisition: G.H., P.S.N.

## MATERIALS AND METHODS

### PDX mouse models

The establishment and characterization of the LuCaP PDX models were described previously (75). PDXs were propagated in vivo in male NOD-scid IL2R-gamma-null (NSG) mice (cat#005557). The collection of tumors for the establishment of PDX lines was approved by the University of Washington Human Subjects Division IRB (IRB #2341). PDX lines were evaluated using histopathology by at least two expert pathologists, and histological phenotypic subtype annotations were orthogonally validated based on transcriptome-derived signature marker expression scores to define phenotypes (4,5,22): adenocarcinoma AR-positive (ARPC), neuroendocrine positive (NEPC), and AR-low, neuroendocrine negative (ARLPC). Resected PDX tumors (300-800 mm^3^) were divided into ∼50mg to ∼100mg pieces and stored at -80°C. Animal studies were approved by the Fred Hutchinson Cancer Research Center (FHCRC) IACUC (protocol 1618) and performed in accordance with the NIH guidelines. For the current study, blood was collected by cardiac puncture from animals bearing PDX tumors (measurable size 300-800 mm^3^).

### Human subjects

UW cohort: Blood samples were collected from men with metastatic castration resistant prostate cancer at the University of Washington (collected under University of Washington Human Subjects Division IRB protocol number CC6932 between years 2014-2021). In this study, 61 plasma samples from 30 patients were analyzed. After initial ultra-low pass whole genome sequencing (ULP-WGS) analysis, 47 plasma samples from 30 patients were retained for further high depth of coverage whole genome sequencing (WGS) analysis. All samples were de-identified prior to ctDNA analysis and we employed a double blinded approach for evaluating clinical phenotype predictions.

DFCI cohort I: Plasma was collected from men diagnosed with mCRPC and treated at the Dana-Farber Cancer Institute (DFCI), Brigham and Women’s Hospital, or Weill Cornell Medicine (WCM) between April 2003 and August 2021. All patients provided written informed consent for research participation and genomic analysis of their biospecimen and blood. The use of samples was approved by the DFCI IRB (#01-045 and 09-171) and WCM (1305013903) IRBs. ULP-WGS data at mean coverage 0.5x (range 0.3x – 0.9x) for 101 patients were published previously (25).

DFCI cohort II: Plasma samples in this cohort were collected from men diagnosed with mCRPC and treated at the Dana-Farber Cancer Institute (DFCI). All patients provided written informed consent for blood collection and the analysis of their clinical and genetic data for research purposes (DFCI Protocol # 01-045 and 11-104). WGS data at mean coverage 27x (range 11x – 44x) (67), and ULP-WGS data at mean coverage 0.13x (range 0.07x – 0.18x) (68, 72) were downloaded from dbGAP accession phs001417. Eleven samples from six patients had matching WGS and ULP-WGS with paired-end reads, necessary for analysis by Griffin. Prostate specific antigen (PSA, ng/mL) values and treatment at the time of the blood draw were previously published (68). The six patients were treated for adenocarcinoma using Abiraterone, Enzalutamide, or Bicalutamide, or the patients had detectable levels of PSA.

Healthy donor plasma cfDNA WGS data used in this study were obtained from previously published studies. Two samples (HD45 and HD46) with coverage of 13x and 15x, respectively, were accessed from dbGAP under accession phs001417 (67, 72). These donors were consented under DFCI protocol IRB (# 03-022). Thirteen healthy donor plasma cfDNA WGS data (12 male: NPH002, 03, 06, 07, 12, 18, 23, 26, 33, 34, 35, 36; 1 female (used in admixtures): NPH004) with coverages between 13.5x – 27.6x were obtained from the European Phenome Archive (EGA) under accession EGAD00001005343 (41).

### PDX plasma processing

Blood samples were collected from NSG mice bearing subcutaneous PDX tumors at the time of sacrifice. The PDX lines were maintained at vivaria in the University of Washington and FHCRC. The blood was processed following methods described for human plasma DNA processing for subsequent DNA isolation. Blood was collected in purple cap EDTA tubes and processed within 4 hours. All blood samples were double spun using centrifugation at 2500g for 10 minutes followed by a 16000g spin of the plasma fraction for 10 minutes at room temperature. For each PDX line, 7-10 mouse plasma samples were pooled. Processed plasma samples were preserved in clean, screw-capped cryo-microfuge tubes and stored at -80°C prior to cfDNA isolation.

### Cell-free DNA isolation

The QIAamp Circulating Nucleic Acid Kit was used to isolate cfDNA from PDX mouse-derived plasma using the recommended protocol. The pooled plasma samples from 7-10 mice for each PDX line contained ∼2-3 mL total plasma volume for each line. The filter retention-based cfDNA kit method does not implement any fragment size class enrichment. Isolated cfDNA was quantified using the Qubit dsDNA HS assay (Invitrogen) and the cfDNA fragment size profiles were analyzed using TapeStation HS D5000 and HS D1000 assays (Agilent).

### Cell-free DNA library preparation and sequencing

For LuCaP PDX mouse plasma samples, NGS libraries were prepared with 50ng input cfDNA. Illumina NGS sequencing libraries were prepared with the KAPA hyperprep kit, adopting nine cycles of amplification, and purified using lab standardized SPRI beads. We used KAPA UDI dual indexed library adapters. Library concentrations were balanced and pooled for multiplexing and sequenced using the Illumina HiSeq 2500 at the Fred Hutch Genomics Shared Resources (200 cycles) and Illumina NovaSeq platform at the Broad Institute Genomics Platform Walkup-Seq Services using S4 flow cells (300 cycles). To match with Illumina HiSeq 2500 data, truncated 200 cycles FASTQ files were generated (100 bp paired end reads).

Clinical patient plasma samples collected at University of Washington (UW cohort) were submitted to the Broad Institute Blood Biopsy Services. Briefly, cfDNA was extracted from 2 mL double-spun plasma and ultra-low-pass whole genome sequencing (ULP-WGS) to approximately 0.2x coverage was performed. The ichorCNA pipeline was used to estimate tumor DNA content (i.e., tumor fraction, see below). Forty-seven samples (from 30 patients) had either ≥ 5% tumor fraction or ≥ 2% tumor fraction with AR amplification observed in ichorCNA and were subsequently sequenced to deeper WGS coverage (∼20x).

### Cell-free DNA sequencing analysis and mouse subtraction

All cfDNA sequencing data used in this study were realigned to the hg38 human reference genome (http://hgdownload.soe.ucsc.edu/goldenPath/hg38/bigZips/hg38.fa.gz). FASTQ files were realigned using BWA (v0.7.17) mem (76). The complete alignment pipeline including configuration settings may be access at https://github.com/GavinHaLab/fastq_to_bam_paired_snakemake.

For PDX ctDNA whole-genome sequence data, we performed mouse genome subtraction following the protocol described previously (77), wherein reads were aligned using BWA mem to a concatenated reference consisting of both human (hg38) and mouse (mm10, GRCm38.p6, http://igenomes.illumina.com.s3-website-us-east-1.amazonaws.com/Mus_musculus/NCBI/GRCm38/Mus_musculus_NCBI_GRCm38.tar.gz) reference genomes. Read pairs where both reads aligned to the human reference genome were retained and all other read pairs were removed. Then, remaining reads were re-aligned to the human-only reference. Finally, the GATK best practices workflow was applied to each sample (78); the complete mouse subtraction pipeline used in this study, including tool versions and parameters, can be accessed at https://github.com/GavinHaLab/PDX_mouseSubtraction. Following mouse subtraction samples with < 3X depth were removed for downstream analysis.

### Cell cycle progression (CCP) score calculation

The 31-gene cell cycle proliferation (CCP) signature (62) was computed from RNAseq data using GSVA (79). The single-sample enrichment scores were calculated with default parameters using genome-wide log2 FPKM values as input for the 31 genes.

### Differential mRNA expression analysis

RNA isolation of 102 tumors from 46 LuCaP PDX samples was performed as described previously (11). RNA concentration, purity, and integrity was assessed by NanoDrop (Thermo Fisher Scientific Inc) and Agilent TapeStation and RNA RIN >=8 was retained for library preparation. RNA-Seq libraries were constructed from 1 ug of total RNA using the Illumina TruSeq Stranded mRNA LT Sample Prep Kit according to the manufacturer’s protocol. Barcoded libraries were pooled and sequenced by Illumina NovaSeq 6000 or Illumina HiSeq 2500 generating 50 bp paired end reads. Sequencing reads were mapped to the hg38 human reference genome and mm10 mouse reference genomes using STAR.v2.7.3a (80). All subsequent analyses were performed in R-4.1.0. Sequences aligning to the mouse genome and therefor derived from potential contamination with mouse tissue were removed from the analysis using XenofilteR (v1.6) (81). Gene level abundance was quantitated using the R package GenomicAlignments v1.32.0 summarizeOverlaps function using mode=IntersectionStrict, restricted to primary aligned reads. We used refSeq gene annotations for transcriptome analysis. Transcript abundances (FPKM) were input to edgeR v3.38.1 (82), filtered for a minimum expression level using the filterByExpr function with default parameters, and then limma v3.52.1 voom was used for differential expression analysis of NEPC vs. ARPC and ARLPC vs. ARPC. We then filtered the results using a list of 1,635 human transcription factors published previously (83), which resulted in 514 genes with FDR<0.05 and fold change > 3. Out of these 514, deregulation of gene expression for 404 transcription factor genes delineated ARPC from NEPC.

### Cleavage Under Targets & Release Using Nuclease (CUT&RUN)

CUT&RUN is an antibody targeted enzyme tethering chromatin profiling assay in which controlled cleavage by micrococcal nuclease releases specific protein-DNA complexes into the supernatant for paired-end DNA sequencing analysis. We performed CUT&RUN assays for three histone modifications, H3K27ac, H3K4me1, and H3K27me3, according to published protocols (46). We performed CUT&RUN on LuCaP PDX tumors using ∼75mg flash-frozen tissue pieces.

Paired-end (50 bp) sequencing was performed and reads were aligned using bowtie2 v2.4.2 (84) to the hg38 human reference assembly. Aligned reads were processed as described in the SEACR protocol (https://github.com/FredHutch/SEACR#preparing-input-bedgraph-files). Peaks were called using SEACR version 1.3 (47) using “stringent” settings and with reference to paired IgG controls. BigWig files were prepared using bamCoverage in deepTools 3.5.0 (85). Genomewide peak heatmap, targeted heatmap, and respective profiles were plotted using deepTools v3.5.0. bigwig formatted files for each phenotype were obtained using the mean function in wiggletools 1.2.8. and deepTools computeMatrix. Phenotype-specific informative region coordinates were obtained from diffBind v3.5.0, and the top 10,000 most significant regions (all with FDR < 0.05) differentially open between ARPC and NEPC lines were used for downstream feature analyses (see Gene body and promoter region selection for additional subsetting criteria applied on a feature-by-feature basis). For heatmaps and profiles the plotHeatmap function was used. We utilized the “Peak Center” option to derive desired heatmaps. These steps were all performed for H3K27ac, H3K4me1 and H3K27me3 antibodies. Scaled heatmap profiles’ area under the curve (AUC) and peak height at the profile center were estimated using deepStats v0.4 (86) (comparable profiles are scaled to 10 units).

### Differential histone post-translational modification (PTM) analysis

Differential PTM analysis was performed with the Diffbind version 2.16.0 package (87) in R-4.0.1 using standard parameters (https://bioconductor.riken.jp/packages/3.0/bioc/html/DiffBind.html). ARPC, NEPC and ARLPC samples were grouped by histopathological and transcriptome signature defined phenotypes described in the “PDX mouse models” section (Supplementary Table S2A). Samples were loaded with the dba function, reads counted with the dba.count function, and contrast specified as phenotype with dba.contrast and a minimum members of 2. Differential peak sites were computed with the dba.analyze function with default settings. Differential peak binding of NEPC and ARLPC was computed against ARPC samples. Unique binding sites in NEPC and ARLPC were catalogued using bedtools v2.29.2 (88). Intergroup differentially bound peaks were annotated using ChIPseeker 1.28.3 (89) and TxDb.Hsapiens.UCSC.hg38.knownGene 3.2.2 in R 4.1.0.

### ATAC-Seq analysis

ATAC-Seq sequence data for 15 tumor samples from 10 PDX lines were published previously (9). These lines included LuCaP PDX lines with ARPC histology (23.1, 77, 78, 81, 96) and NEPC histology (two replicates each of 49, 93, 145.1, 173.1 and one replicate of 145.2). Paired end reads were aligned using bowtie2 v2.4.2 (84) to the UCSC hg38 human reference assembly with the “very-sensitive” “-k 10” settings. Peaks were called using Genrich version 0.6.1 (https://github.com/jsh58/Genrich). Differential binding analysis was performed using Diffbind version 3.5.0 package in R version 4.1.0. ENCODE blacklisted regions were excluded using hg38- blacklist.v2 (90) (https://github.com/Boyle-Lab/Blacklist). Phenotype specific binding sites were isolated by first selecting for positive fold change open chromatin enrichment and then using Intervene 0.6.5 (91) where regions were considered overlapping if they shared at least 1 bp. Regions with FDR adjusted p-values < 0.05 were then subset to those overlapping the 338,000 established TFBSs (338 TFs x 1,000 binding sites, see Griffin analysis for site selection) by at least 1 bp using BedTools v2.30.0 Intersect. Only regions that overlapped an established TFBS were retained.

### Nucleosome profiling of ctDNA

Griffin is a method for profiling nucleosome protection and accessibility on predefined genomic loci (49). Griffin filters sites by mappability, estimates and corrects GC bias on a per fragment level, and generates GC-corrected coverage profiles around each site. First, griffin takes a site list and examines the mappability in a window (+/- 5000 bp around each site). Mappability (hg38 Umap multi-read mappability for 50bp reads) was obtained from UCSC genome browser (92) (https://hgdownload.soe.ucsc.edu/gbdb/hg38/hoffmanMappability/k50.Umap.MultiTrackMappabi <u>lity.bw</u>). Sites with <0.95 mappability were excluded from further analysis. Next, GC bias was quantified for each sample using a modified version of the approach described previously (93). Briefly, for each possible fragment length and GC content, the number of reads in a bam file and the number of genomic positions with that specific length and GC content were counted. The GC bias for each fragment length and GC content was calculated by dividing the number of observed reads by the number of observed genomic positions for that fragment length and GC content. The GC bias for all possible GC contents at a given fragment length was then normalized to a mean bias of 1. GC biases were then smoothed by taking the median of values for fragments with similar lengths and GC contents (k nearest neighbors smoothing) to generate smoothed GC bias values.

After GC correction, nucleosome profiling was performed in each sample. For each mappable site of interest, fragments aligning to the region ± 5000 bp from the site were fetched from the bam file. Fragments were filtered to remove duplicates and low-quality alignments (<20 mapping quality) and by fragment length. Nucleosome size fragments (140-250 bp) were retained. Fragments were then GC corrected by assigning each fragment a weight of 1/GC_bias for that given fragment length and GC content and the fragment midpoint was identified. The number of weighted fragment midpoints in 15bp bins across the site were counted. For composite sites, all sites of a given type (such as all sites for a given transcription factor) were summed together to generate a single coverage profile. Individual or composite coverage profiles were normalized to a mean coverage of 1 in the ± 5000bp region surrounding the site. Finally, sites were smoothed using a Savitsky-Golay filter with a window length of 165bp and a polynomial order of 3. The window ± 1000 bp around the site was retained for plotting and feature extraction (See Griffin manuscript for further details); when plotting sites, shading illustrates the 95% confidence interval within sample groups. Features extracted from individual or composite sites included:

a) “mean central coverage,” the mean coverage between -30 to 30 bp relative to the site center,
b) “mean window coverage,” the mean coverage between -990 to 990 bp relative to the site center, and
c) “max wave height,” the absolute difference between the minimum coverage within the window from -120 to 30 bp and maximum coverage in the window from 31 to 195 bp relative to the TSS.

### Analysis of selective transcription factor binding sites (TFBS)

Transcription factor binding site (TFBS) Griffin analysis was conducted with the same TFBS list utilized in Griffin (49). After intersecting these 338 with 404 differentially expressed TFs identified through RNA-Seq 107 remained, on which we performed unsupervised hierarchical clustering of central window mean values (see Griffin analysis). Hierarchical clustering was performed using the Ward.D2 method with Euclidean distance and complete linkage settings; the groupings were determined using cutree_cols=2 for columns (LuCaP CRPC phenotypes) and cutree_rows=13 for rows (TFs) on the dendrograms.

### Gene body and promoter region selection

For individual gene body and promoter analyses Ensembl BioMart v104 (hg38) (94) was used to directly retrieve protein coding transcript start (TSS) and end (TES) coordinates. For promoter region analysis the window ±1000 bp relative to the TSS was considered. For gene body analysis, the region between the TSS and TES was considered. In the case of genes with multiple transcripts, analyses were limited to the longest transcript resulting in 19,336 regions. In downstream analysis of LuCaP PDX cfDNA, if any lines did not meet specific criteria in a region (including differentially open histone modification regions) that feature/region combination was excluded from analysis, leading to a variable lower number of regions considered based on the feature. These criteria included requiring at least 10 total fragments in a region for all Fragment size analysis (see below) and a non-zero number of “short” and “long” fragments for the short-long ratio; short-long ratios less than 0.01 or greater than 10.0 were also excluded as outliers. For Phasing analysis (see below) we also excluded amplitude components and thus NPS where individual components were 0, or where the ratio was less than 0.01 or greater than 10.0, indicative of insufficient coverage. In the case of mean phased nucleosome distance, if no peaks were identified or the value in a region exceeded 500 (indicative of highly irregular/sparse pileups also from low coverage) those regions were also excluded. Any region with no coverage in a line was excluded from all analyses. This resulted in gene lists that differed in numbers between genomic contexts and feature types.

### Fragment size analysis

Fragments were first filtered to remove duplicates and low-quality alignments (<20 mapping quality) and by fragment length (15-500 bp). In individual genomic loci/windows, we computed the fragment short-long ratio (FSLR) as the ratio of short (15 - 120 bp) to long (140 - 250 bp) fragments. We also calculated the mean, median absolute deviation (MAD: median(|*X_i_* median(*X*)|)), and coefficient of variation (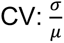 where σ = standard deviation, μ = mean) of the fragment length distribution for each selected window. The fragment size analysis code and implementation used in this study can be accessed at https://github.com/GavinHaLab/CRPCSubtypingPaper/tree/main/FragmentAnalysis.

### Nucleosome phasing analysis (TritonNP)

Fragments were first filtered to remove duplicates and low-quality alignments (<20 mapping quality) and by fragment length (nucleosome-sized: 140-250 bp). Next we performed fragment-level GC bias correction utilizing the same pre-processing method defined in Griffin. A band-pass filter was then applied to the corrected coverage in each region of interest by taking the Fast Fourier Transform (FFT) (scipy.fft v1.8.0) (95) and removing high-frequency components corresponding to frequency components < 146 bp before reconstructing the signal. This cutoff was chosen to ensure that periodic fit signal for downstream evaluation must come from the minimum possible inter-nucleosome distance, thus excluding peak pileups that would not indicate an overall trend in nucleosome phasing. Local peak calling was then done on the smoothed signal to infer average inter nucleosome distance or “phased nucleosome distance” by finding maxima directly. To quantify clarity of overall phasing we took the average frequency amplitude in two bands corresponding to a core + linker (180-210 bp) and core only (150-180 bp), with the former measuring the strength of typically aligned nucleosomes and the latter giving a measure of the underlying signal strength not coming from either high frequency noise or low frequency shifts in total coverage. The ratio of these two amplitude averages forms the Nucleosome Phasing Score (NPS). Because peak locations are assumed to be independent of copy number alterations or depth, and the NPS by virtue of being a ratio divides out any confounding DNA/depth variation between sites, both features are taken as agnostic of CNAs or variable depth. Code and implementation of the method can be found at https://github.com/denniepatton/TritonNP.

### ctDNA tumor-normal admixtures and benchmarking

Admixtures for evaluating benchmarking performance were constructed using 5 ARPC (LuCaP 35, 35CR, 58, 92, 136CR) and 5 NEPC (LuCaP 49, 93, 145.2, 173.1, 208.4) lines mixed to 1%, 5%, 10%, 20%, and 30% tumor fraction with a single healthy donor plasma line (NPH004, EGAD00001005343) at ∼25X mean coverage, assuming 100% tumor fraction in post-mouse subtracted PDX sequencing data. After extracting chromosomal DNA with SAMtools v1.14 (96) and removing duplicates with Picard (https://broadinstitute.github.io/picard/), SAMtools was used to merge BAM files. Admixtures were then down-sampled to the number of reads corresponding to 1X and 0.2X using SAMtools to evaluate (ultra) low-pass WGS performance. During unsupervised benchmarking of each admixture the healthy and LuCaP line used in the admixture were excluded from the generation of feature distributions to ensure the model would not learn from the lines being interrogated. The admixture pipeline used in this study can be accessed at https://github.com/GavinHaLab/Admixtures_snakemake.

### Supervised binary classification of ARPC and NEPC

Binary classification of ARPC and NEPC subtypes using individual region and feature combinations was conducted using XGBoost v1.4.2 ‘XGBClassifier’ implemented in Python with default parameters. Features included NPS and Mean Phased Nucleosome Distance (see Phasing analysis) in histone modification regions, promoters, and gene bodies; fragment size mean, short-long ratio, and coefficient of variation (see Fragment size analysis) in histone modification regions, promoters, and gene bodies; central and window coverage (see Griffin analysis) in promoters, composite TFBSs, and composite differentially open chromatin regions identified through ATAC-Seq; and Max Wave Height (See Griffin analysis) in promoters. We applied stratified 6-fold cross-validation where two ARPC samples and one NEPC sample was held out in each fold. This was repeated 100 times and performance was computed using area under the receiver operating characteristic (ROC) curve (AUC) and 95% confidence intervals for each individual feature and region combination. Code and implementation of the method can be found at https://github.com/GavinHaLab/CRPCSubtypingPaper/tree/main/SupervisedLearning.

### Tumor fraction estimation

Tumor fractions from patient plasma samples were assessed using ichorCNA (72) with binSize 1,000,000 bp and hg19 reference genome. Default tumor fraction estimates reported by ichorCNA were used. See https://github.com/GavinHaLab/CRPCSubtypingPaper/tree/main/ichorCNA_configuration for complete configuration settings.

### Phenotype prediction model (ctdPheno)

We developed a probabilistic model to classify the mCRPC phenotype (ARPC or NEPC) in an individual patient plasma ctDNA sample. This is a generative mixture model that is unsupervised—it does not train on the patient cohort of interest. However, the model accepts the pre-estimated tumor fraction from ichorCNA for the given patient ctDNA sample, as well as the pre-computed ctDNA features values from the LuCaP PDX ctDNA and healthy donor ctDNA as prior information. For each patient ctDNA sample, it fits the heterogeneous tumor fractions against the pure PDX LuCaP models. The expected feature value (mean m and standard deviation s) from each phenotype *k* for feature *i* were taken from the mean of LuCaP PDX samples (*μ*_*i,k*_), or taken from the mean of a panel of normals *H* (*μ*_*i,H*_, male only, n = 14; see Healthy Donor cohort) assuming a Gaussian distribution, is shifted such that the shifted values m’i,k , s’i,k took the form:

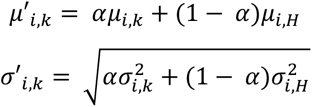

where a is the tumor fraction estimate for each test sample. In the final model, four features were used: composite open chromatin regions (central and window mean coverage) for specific phenotypes (ARPC and NEPC) identified from the LuCaP PDX ATAC-Seq analysis using Griffin (see Griffin analysis). For each feature *i*, we then found the probability that the observed sample came from a mixture of the tumor-fraction-corrected Gaussian distributions, where *θ* is the NEPC mixture weight:

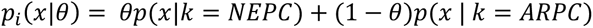

The *θ* parameter is estimated by maximizing the joint log-likelihood *L* for a given patient sample:

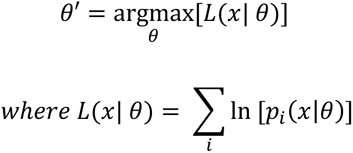

*θ* has range [0,1], where higher values indicate an increased proportion of the sample having a NEPC phenotype and was used as the NEPC prediction score metric. Code and implementation of the method can be found at https://github.com/GavinHaLab/CRPCSubtypingPaper/tree/main/ctdPheno.

### Analysis and classification of clinical patient samples

After establishing feature distributions using the LuCaP PDX lines and normal panel as described in Generative model, the model was applied to three clinical patient cohorts (see Human subjects for cohort information). Initial scoring using the model was run on DFCI cohort I, consisting of 101 ULP-WGS samples with paired-end reads. Tumor fraction estimates predicted by ichorCNA and tumor phenotype classifications were obtained from the original study (25). A prediction score threshold of 0.3314 for calling NEPC was chosen because it offered an optimal performance for sensitivity (90%) and specificity (97.5%), where sensitivity is the true positive rate for identifying

NEPC samples 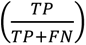 and specificity is the true negative rate for identifying ARPC samples 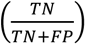. Alternative thresholds maximizing sensitivity and specificity were 0.1077, at which 95% sensitivity was achieved with a lower specificity of 93.8%, and 0.3769 with a lower sensitivity of 81.0% but higher specificity of 98.8%. To compare these predictions with cfDNA methylation (cfMeDIP-seq) classification on the same plasma samples in DFCI cohort I, the concordance was computed between the ctdPheno NEPC prediction score and the cfMeDIP NEPC score obtained from the original study using a 0.15 threshold (25).

We then validated the model on two cohorts, beginning with the already published DFCI cohort II (67,68,72). We restricted our analysis to eleven samples from six patients with matched ULP-WGS and WGS data with paired-end reads. Tumor fraction estimates from ichorCNA were obtained from the original study (72). All samples were considered adenocarcinoma (ARPC) based on clinical histories (see Human subjects). The scoring threshold of 0.3314, determined from DFCI cohort I was used for phenotype classification.

For the *UW cohort*, consisting of 47 samples from 30 patients, ichorCNA was used to estimate sample tumor fractions as described above, while clinical phenotype was determined from clinical histories and expert chart review. We evaluated model performance on matched ULP-WGS and WGS data for unambiguous clinical phenotypes of ARPC and NEPC. The chosen scoring threshold of 0.3314 was used, and the fraction of correctly predicted ARPC (n=26) and NEPC (n=5) was computed. The remaining 16 samples with mixed histologies were not evaluated for performance.

## STATISTICAL ANALYSIS

Quantification of and statistical approaches for high-throughput sequencing data analysis are described in the methods above. When non-parametric distributions (not normally distributed) of numerical values of a particular parameter in a population were compared (using boxplots or in tables), the two-tailed Mann-Whitney U test (also known as the Wilcoxon Rank Sum test; scipy.stats.mannwhitneyu, (95) was used to test if any two distributions being compared were significantly different, with Benjamini-Hochberg (statsmodels.stats.multitest.fdrcorrection, https://www.statsmodels.org) correction applied in multiple testing scenarios. All boxplots represent the median with a centerline, interquartile range (IQR) with a box, and first quartile – 1.5 IQR and third quartile + 1.5 IQR with whiskers. PCA was conducted in Python (sklearn.decomposition.PCA; https://scikit-learn.org)

## DATA AVAILABILITY

LuCaP patient derived xenograft (PDX) sequencing data generated in this study, including CUT&RUN results and processed cfDNA (cfDNA) sequencing data will be deposited at GEO and will be publicly available as of the date of publication. LuCaP PDX plasma cell-free DNA whole genome sequencing data will be deposited in dbGaP. The patient plasma genome sequencing data generated in this study will be deposited in a public repository and will be publicly available as of the date of publication. This paper also analyzes existing, publicly available data, including LuCaP PDX RNA-Seq (GSE199596) and ATAC-Seq data (SE156292). The DOIs and links to specific tools are available in the methods.

Any additional information required to reanalyze the data reported in this paper is available from the lead contact upon request.

