## Supplementary Data for "Nucleosome patterns in circulating tumor DNA reveal transcriptional regulation of advanced prostate cancer phenotypes"

* These authors contributed equally.

† Co-senior authors.

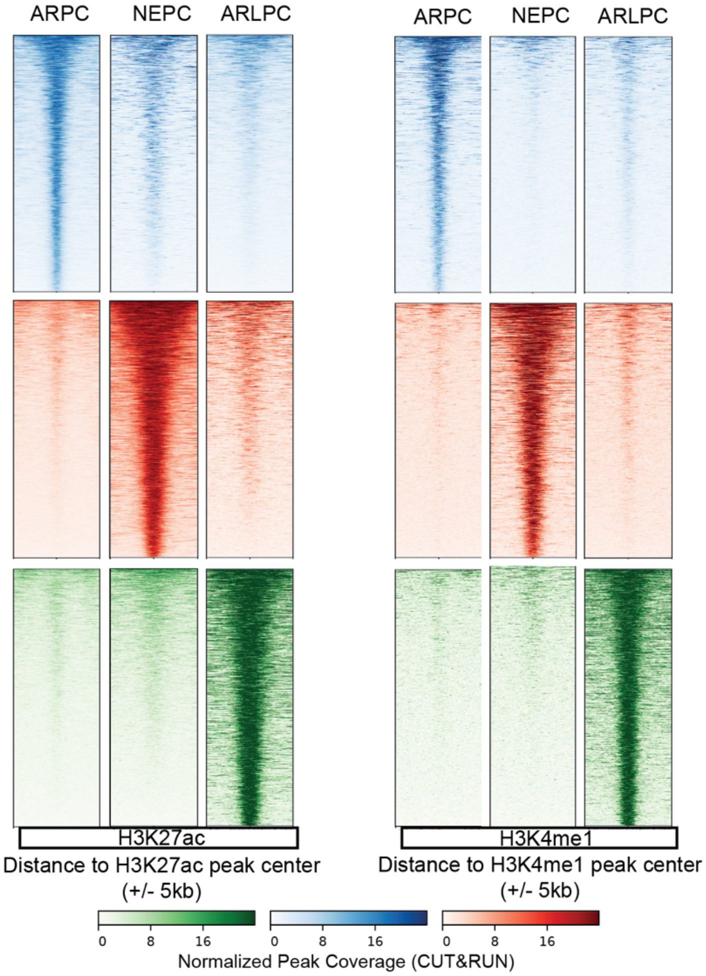


**Supplementary Fig. S1**

Differential global histone tail modification heatmap analysis between androgen regulated prostate cancer (ARPC: blue), neuroendocrine prostate cancer (NEPC: red) and AR low activity prostate cancer (ARLPC: green) suggesting a significant number of lineage associated characteristic histone tail PTM peaks. Normalized H3K27ac and H3K4me1 Heatmap showing peak densities at differential PTM loci (+/- 5kb from peak center) between respective lineage subtypes.

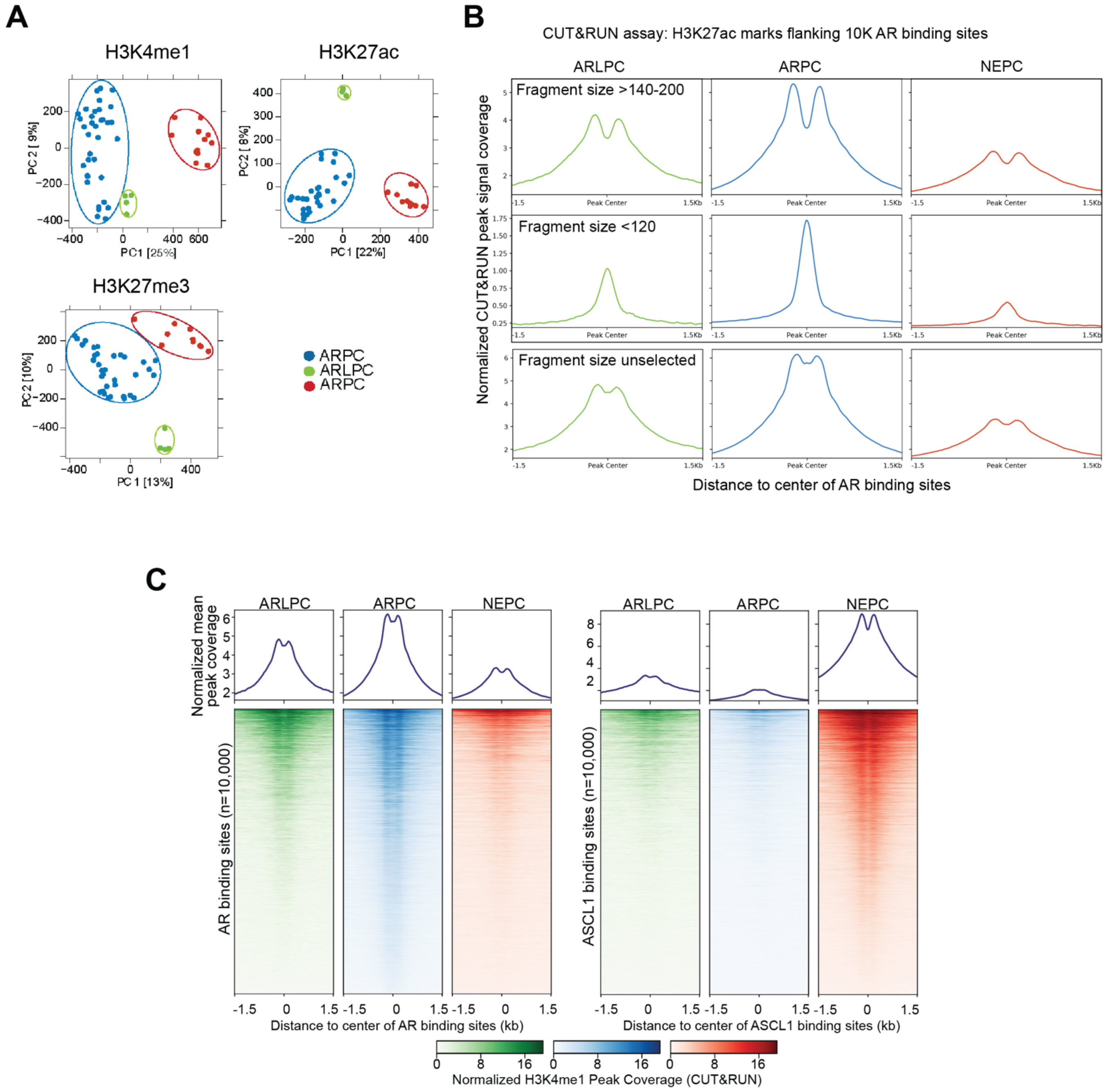


**Supplementary Fig. S2**

**(A)** PCAs of global histone post translational modification (PTM) marks on up to 33 LuCaP patient derived xenografts (PDXs) suggesting distinctive subtype clustering based on tumors. From left to right, all SEACR pipeline called H3K4me1, H3K27ac, and H3K27me3 peaks are considered for respective PCAs.

**(B)** The H3K27ac normalized relative intensity profile suggests AR binding sites in ARPC have a deep nucleosome depleted region (NDR) with large nucleosome fragment sizes selected (140-200nt) (**top panel**). The NDR dip feature is compromised with sub nucleosome fragment sizes (<120nt) (**mid panel**) or without fragment size selection (**bottom panel**). In ARLPC or NEPC, the overall profile signal is expectedly low. The androgen receptor binding sites (ARBS) NDR feature ARLPC or NEPC is similarly impacted by the fragment size of the CUT&RUN data.

**(C)** SEACR called H3K4me1 peaks of key mCRPC subtypes ARLPC, ARPC, and NEPC flanking either AR (left) or ASCL1 (right) binding sites identify differential open chromatin status along the key subtype marker transcription factor (TF). The analysis is limited to CUT&RUN derived 140-200 nt fragments. The 10,000 sites composite peak profile in ARPC suggests a relatively higher frequency of accessible nucleosomes flanking to the chosen AR binding sites suggestive of higher transcription activity. The 10,000 sites composite peak profile in NEPC flanking nucleosome are more frequently accessible in NEPC.

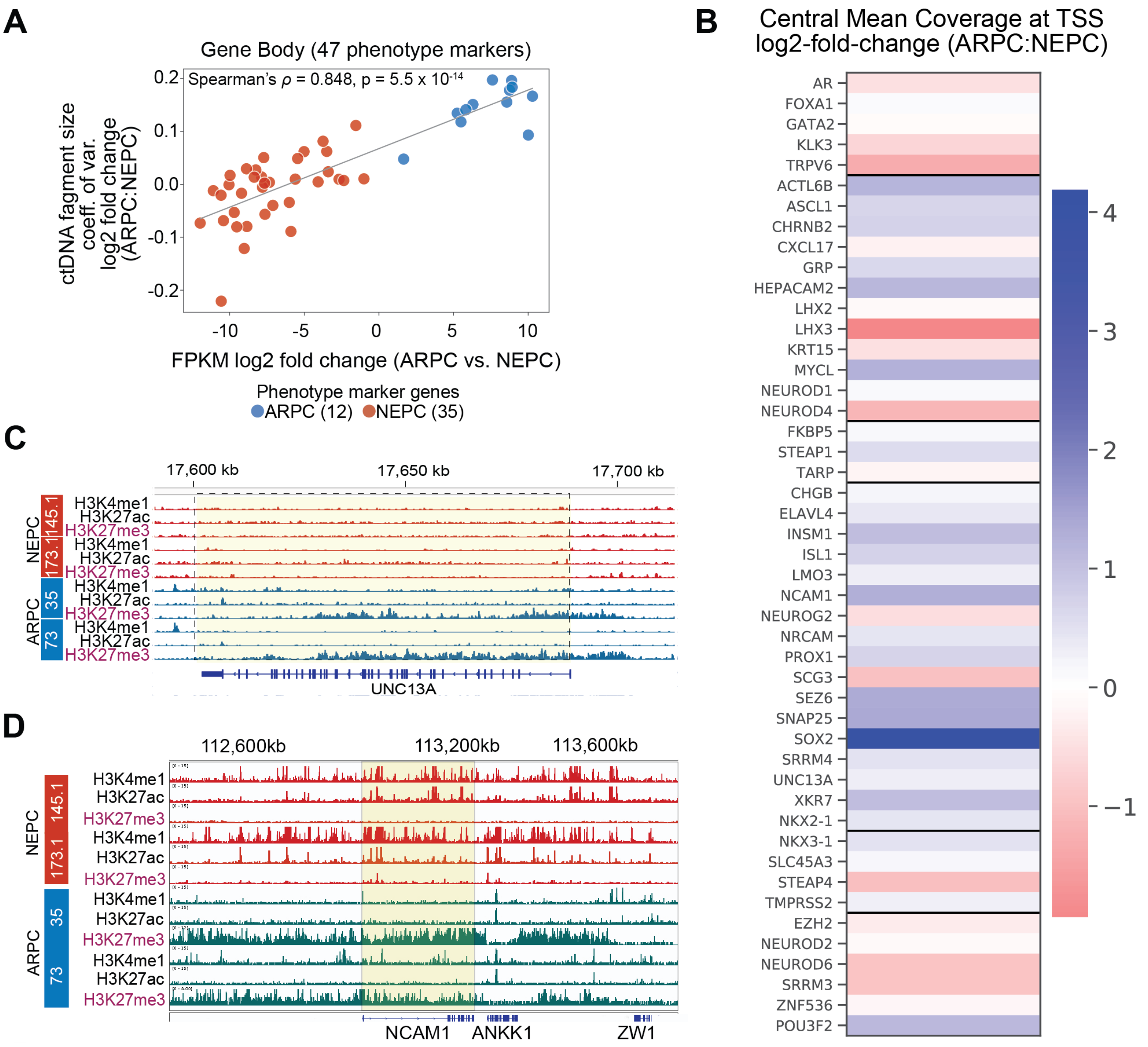


**Supplementary Fig. S3**

**(A)** Heatmap of Griffin mean central coverage Log2 fold change between ARPC and NEPC lines for 47 phenotype defining genes' promoter regions (+/- 1,000 bp around the TSS).

**(B)** CUT&RUN coverage at post-translational modifications (PTM) for UNC13A in select ARPC and NEPC lines illustrates gene body H3K27me3 in ARPC lines lead to activity repression in the absence of activation marks (H3K27ac, H3K4me1).

**(C)** CUT&RUN coverage at post-translational modifications (PTM) for NCAM1 in select ARPC and NEPC lines illustrates H3K27me3 activity in ARPC lines lead to repression in the absence of activation marks (H3K27ac, H3K4me1).

**(D)** Comparison of the $\log_2$ fold change (ARPC/NEPC) of mean mRNA expression vs mean coefficient of variation (CV) in the 47 phenotypic lineage marker genes' promoter regions.

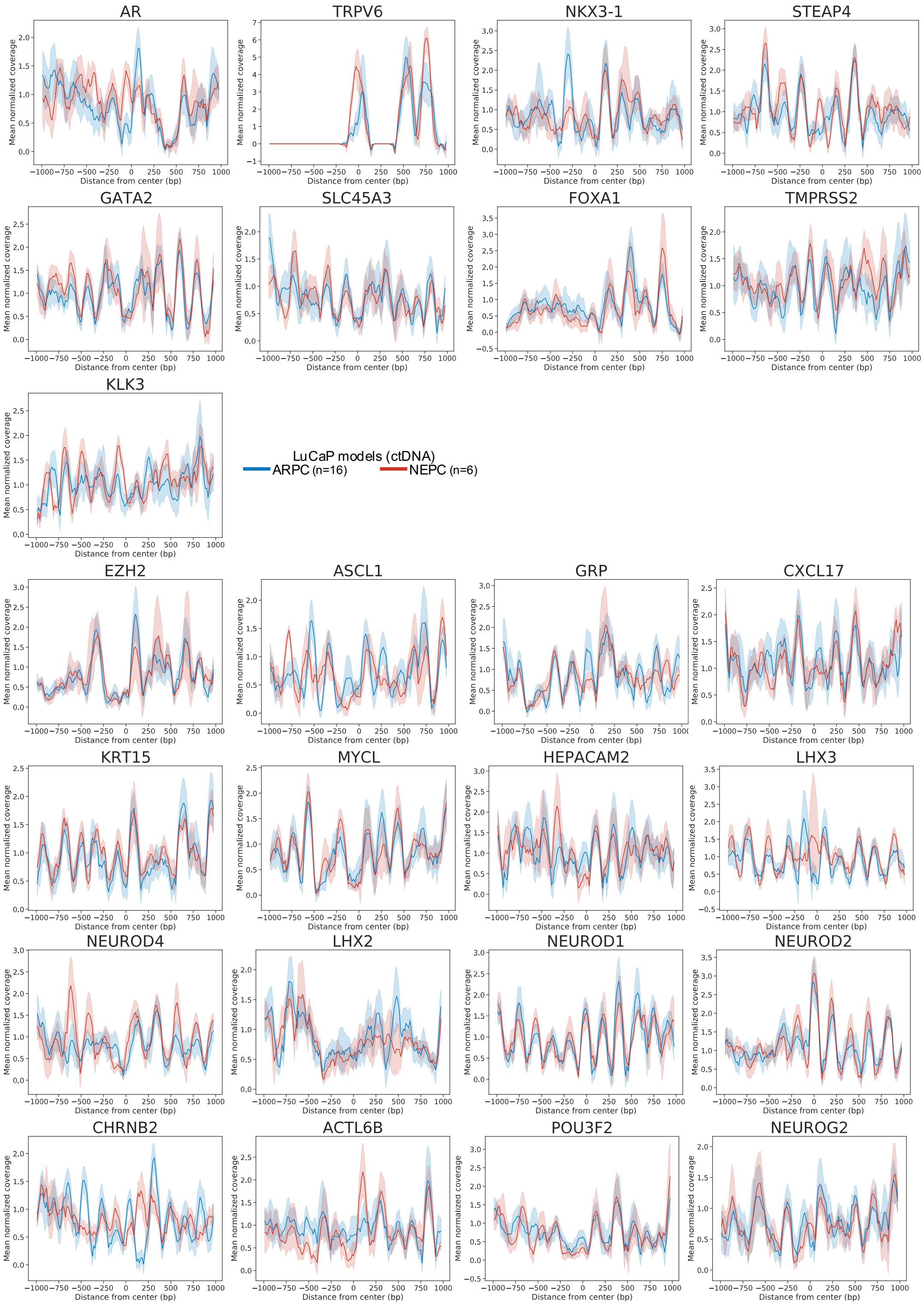


**Supplementary Fig. S4**

Coverage profiles at TSS binding sites in ctDNA analyzed using Griffin for genes in group 1 (Figure 2D). Coverage profile means (lines) and 95% confidence interval with 1000 bootstraps (shading) are shown.

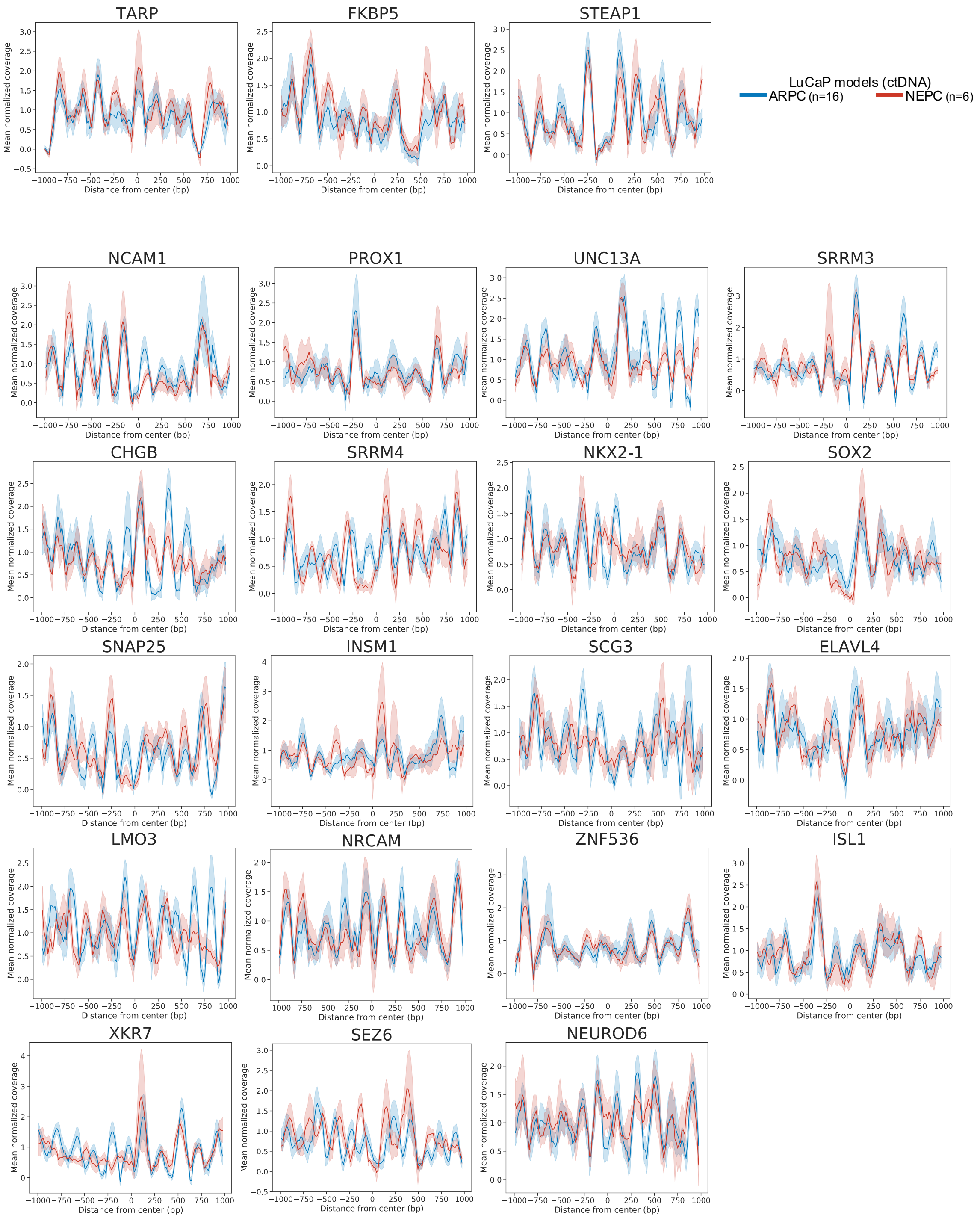


**Supplementary Fig. S5**

Coverage profiles at 1000 TSS binding sites in ctDNA analyzed using Griffin for genes in group 2 (Figure 2D). Coverage profile means (lines) and 95% confidence interval with 1000 bootstraps (shading) are shown.

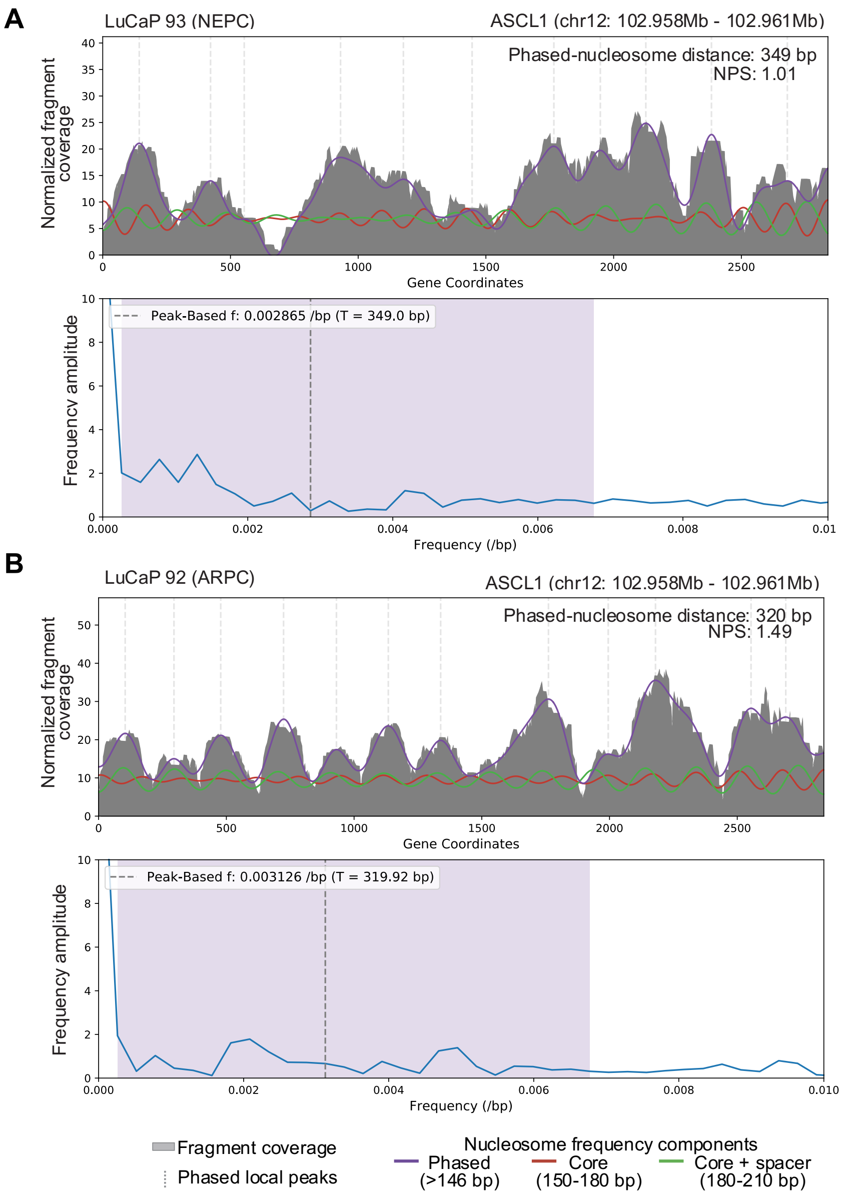


**Supplementary Fig. S6**

**(A)** (**top**) illustration of nucleosome phasing feature extraction for the subtype-defining gene ASCL1 shows disordered phasing in NEPC PDX line LuCaP 93; a band-pass filter-based smoothing method is used to isolate peaks originating from nucleosome-sized read pileups in GC-corrected fragment coverage, and peak calling is used to infer mean inter nucleosome distance. NPS = 1.01. (**bottom**) frequency space of the same gene body with the frequency band corresponding to the purple fit line highlighted (constant value at 0 /bp is included in the upper profile for visualization but does not impact peak calling); the dotted line shows where the mean phased-nucleosome distance falls in the spectrum.

**(B)** (**top**) illustration of nucleosome phasing feature extraction for the subtype-defining gene ASCL1 shows ordered phasing in NEPC PDX line LuCaP 92, with a higher NPS score of 1.49 and smaller phased-nucleosome distance. (**bottom**) frequency space illustrates a shift towards higher frequenies (larger periodic signal) compared to (A).

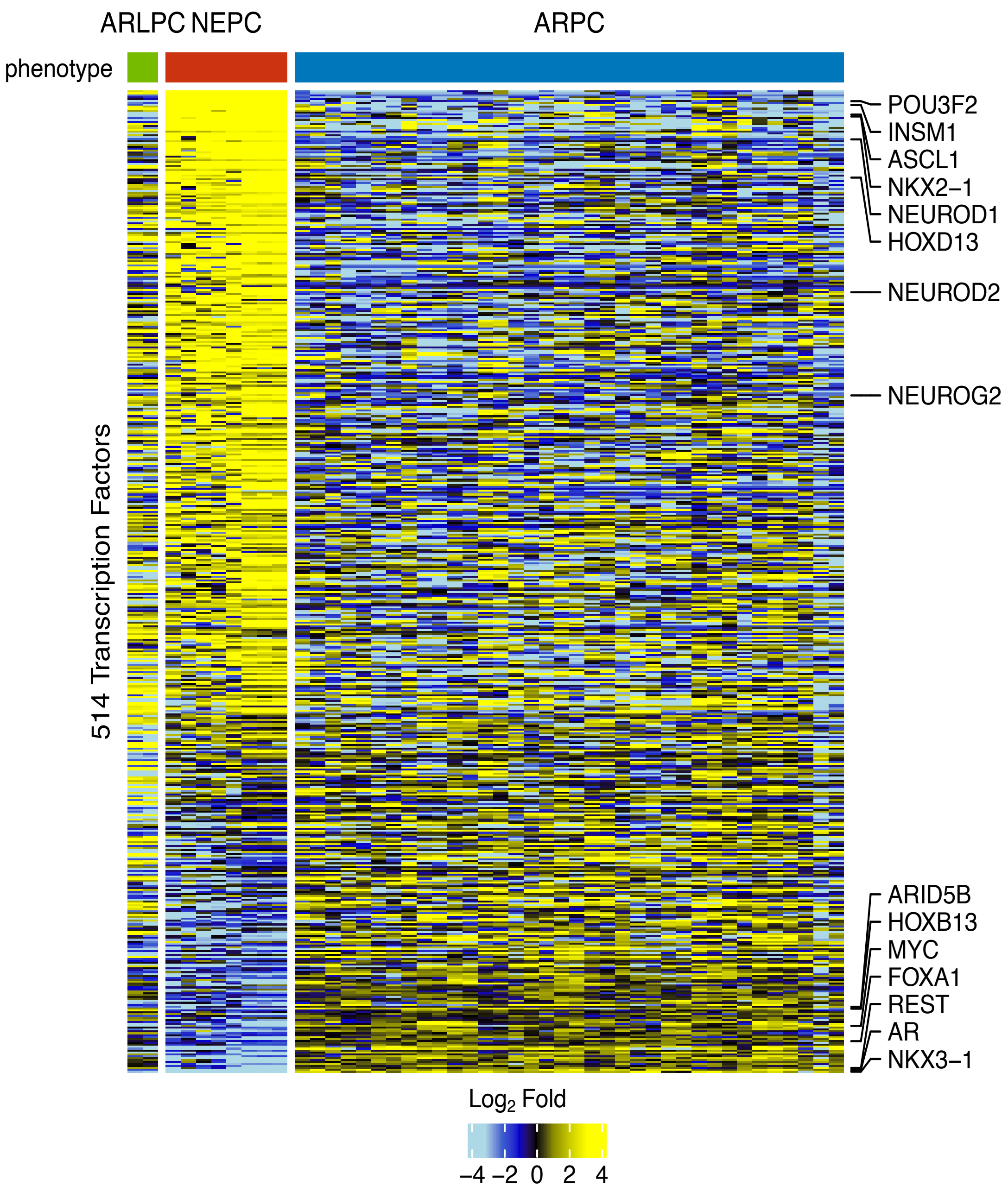


**Supplementary Fig. S7**

RNA-Seq heatmap of top differentially expressed transcription factor genes in 102 tumors from 46 LuCaP PDX lines with well-defined phenotypic subtype annotation (ARPC, NEPC, ARlowPC). Mean $\log_2$ fold change per LuCaP line is shown for 514 transcription factor genes with FDR < 0.05 and fold change > 3.

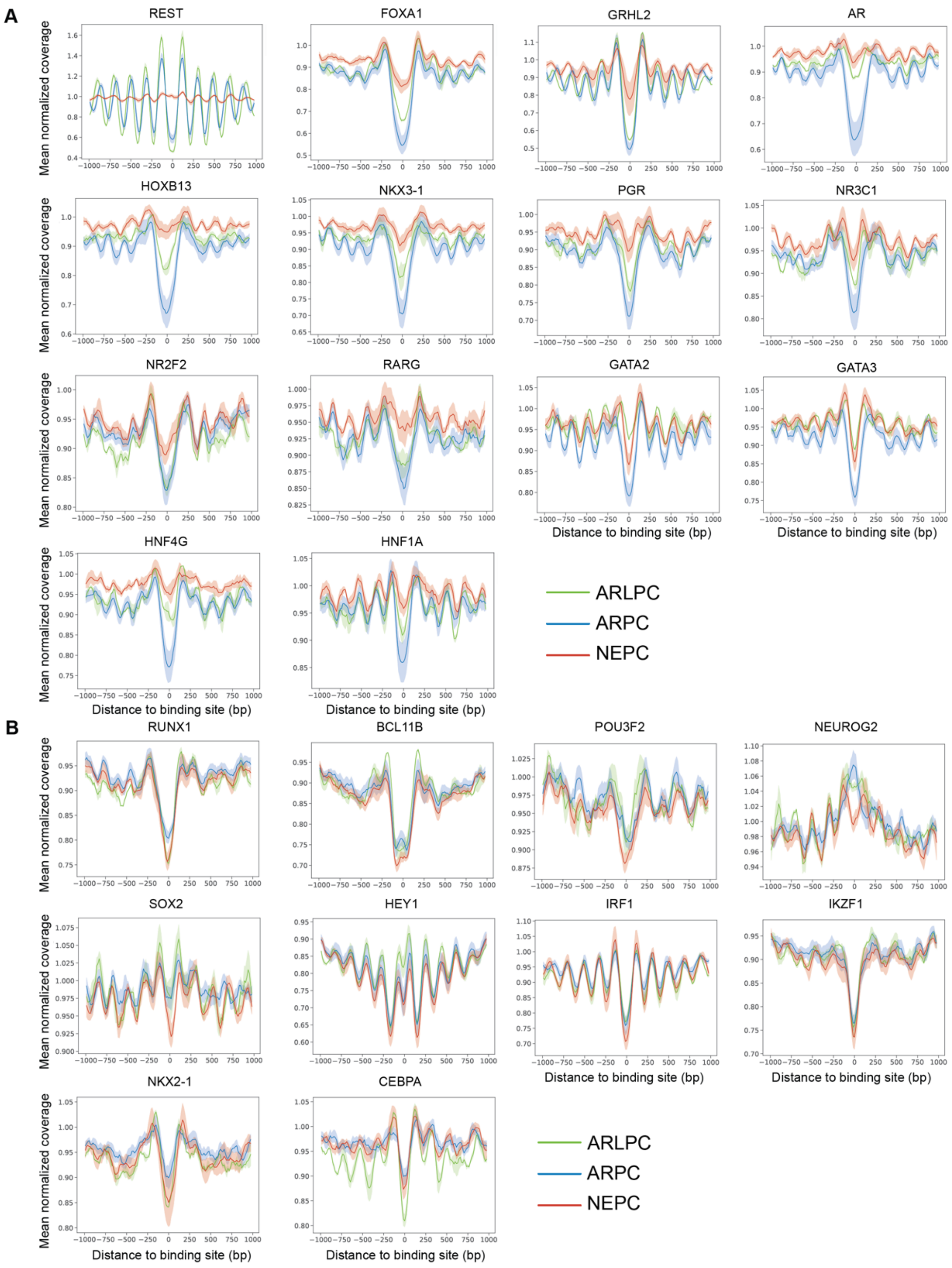


**Supplementary Fig. S8**

**(A)** Griffin coverage profiles for ARPC, NEPC, and ARLPC PDX lines in 14 TFs. In mRNA expression analysis, the selected TFs are significantly upregulated in ARPC relative to NEPC.

**(B)** Griffin coverage profiles for ARPC, NEPC, and ARLPC PDX lines in 10 TFs. In mRNA expression analysis, the selected TFs are significantly downregulated in ARPC relative to NEPC. To note: In these selected downregulated TF we did not notice a significant difference in Griffin central coverage or window mean coverage between 3 phenotypic subtypes of prostate cancer (ARPC, NEPC and ARLPC).

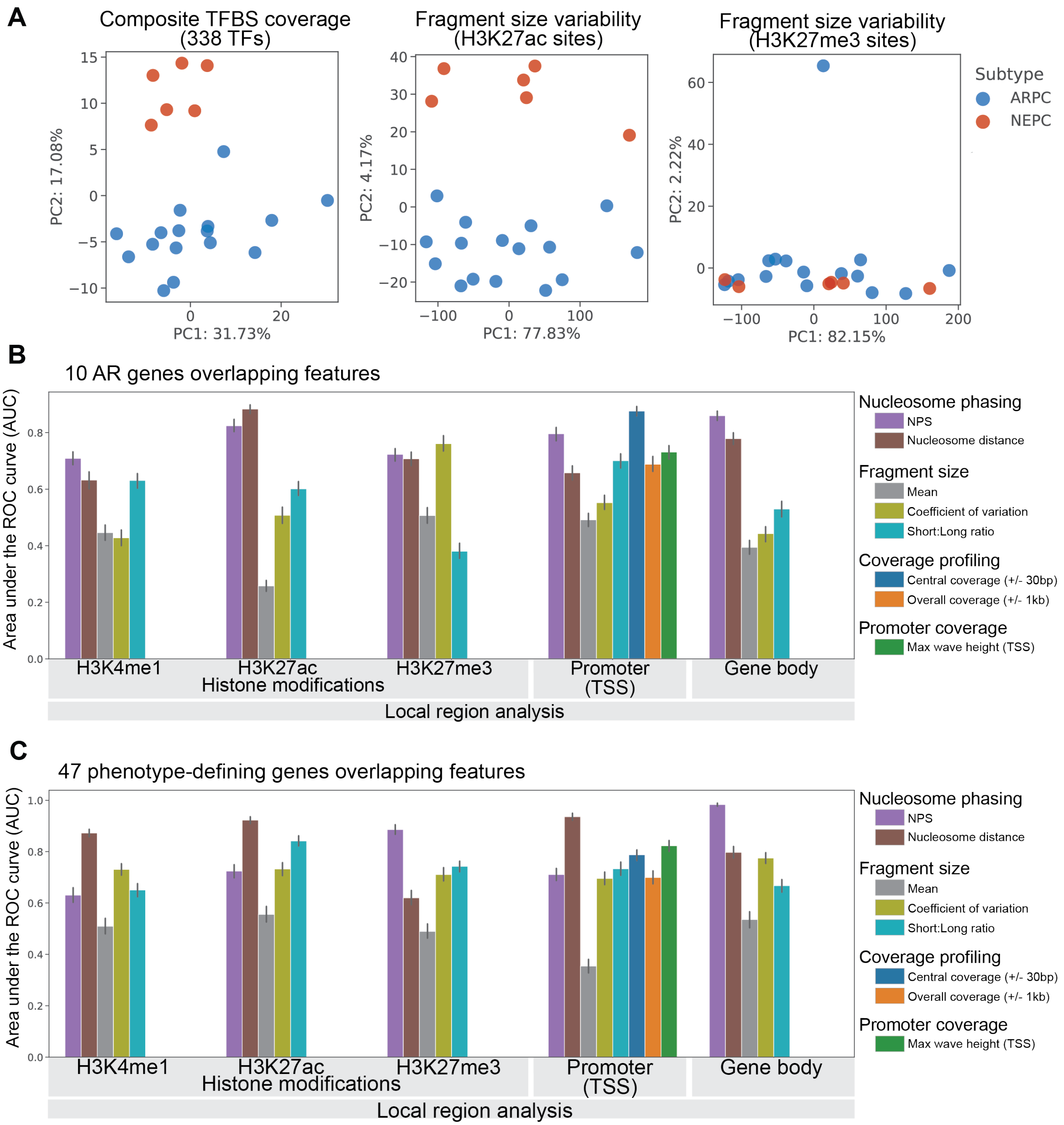


**Supplementary Fig. S9**

**(A)** PCA plots of ARPC and NEPC PDX lines for feature and region combinations: (**left**) Griffin central coverage means for all 338 queried TFBSs; (**center**) fragment-size variation (CV) in global H3K27ac sites (n = 9,689); (**right**) fragment-size variation (CV) in global H3K27me3 sites (n = 9,655).

**(B)** Bar plot of AUCs for subtyping ARPC vs NEPC PDX lines using 100 repeats of stratified 6-fold cross validation with supervised machine learning (XGBoost) in ‘ARG10’ overlapping regions.

**(C)** Bar plot of AUCs for subtyping ARPC vs NEPC PDX lines using 100 repeats of stratified 6-fold cross validation with supervised machine learning (XGBoost) in the ‘phenotype 47’ overlapping regions.

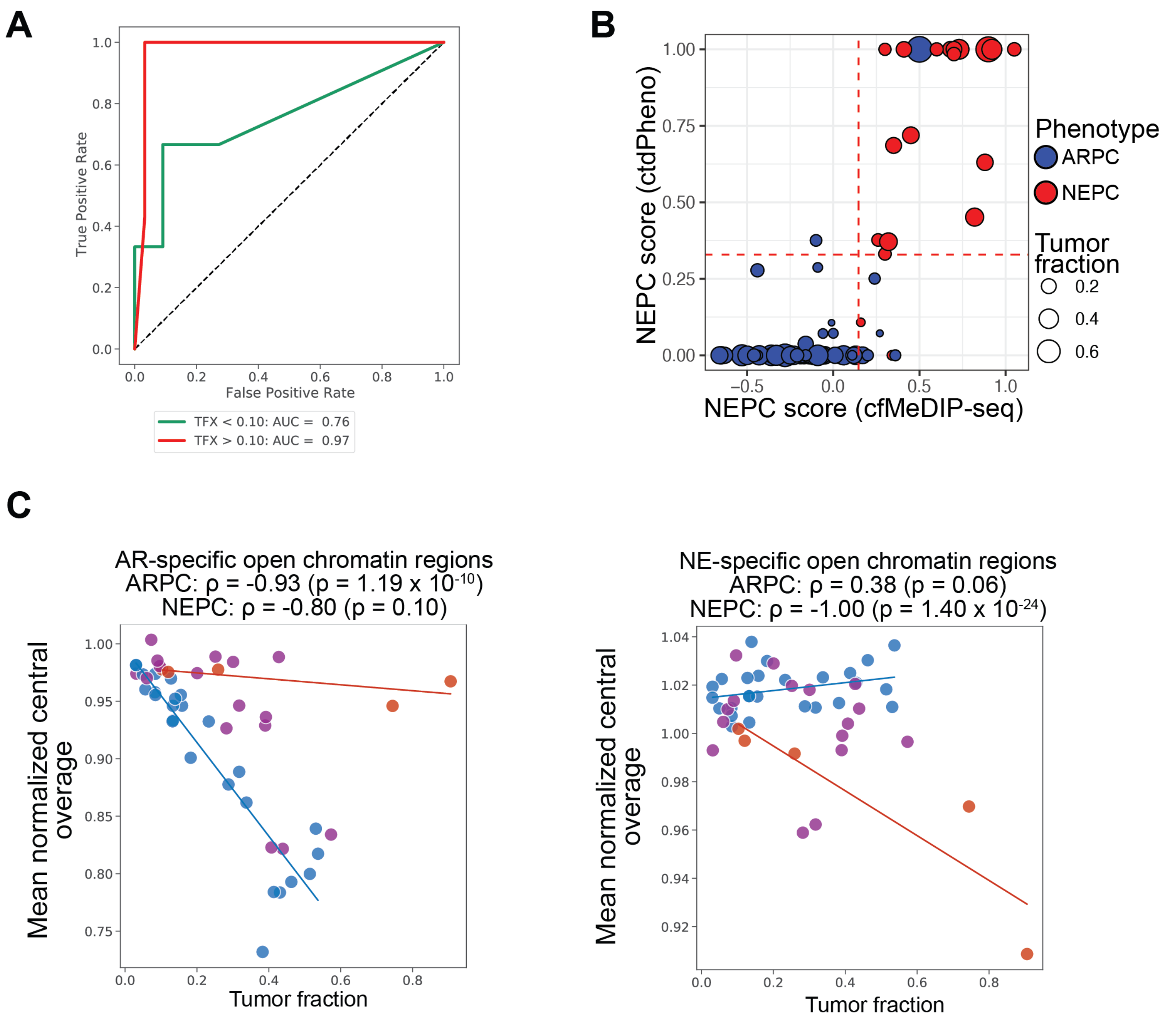


**Supplementary Fig. S10**

**(A)** ROC of the unsupervised generative model for 101 ULP-WGS of mCRPC patients (80 ARPC, 21 NEPC; DFCI cohort I) divided by tumor fraction < 0.10 and >= 0.10, with AUC of 0.97 and 0.76, respectively.

**(B)** Comparison of NEPC scores between ctdPheno and cfMeDIP-seq on the same plasma samples for the DFCI cohort I (N=101).

**(C)** (**left**) Composite ARPC open chromatin region central means vs tumor fraction for 47 WGS patient samples (UW cohort) colored by histology; fit lines through ARPC samples and NEPC samples with corresponding Spearman correlation coefficient and p-values. (**right**) Composite NEPC open chromatin region central means vs tumor fraction for 47 WGS patient samples colored by histology; fit lines through ARPC samples and NEPC samples with corresponding Spearman correlation coefficient and p-values.

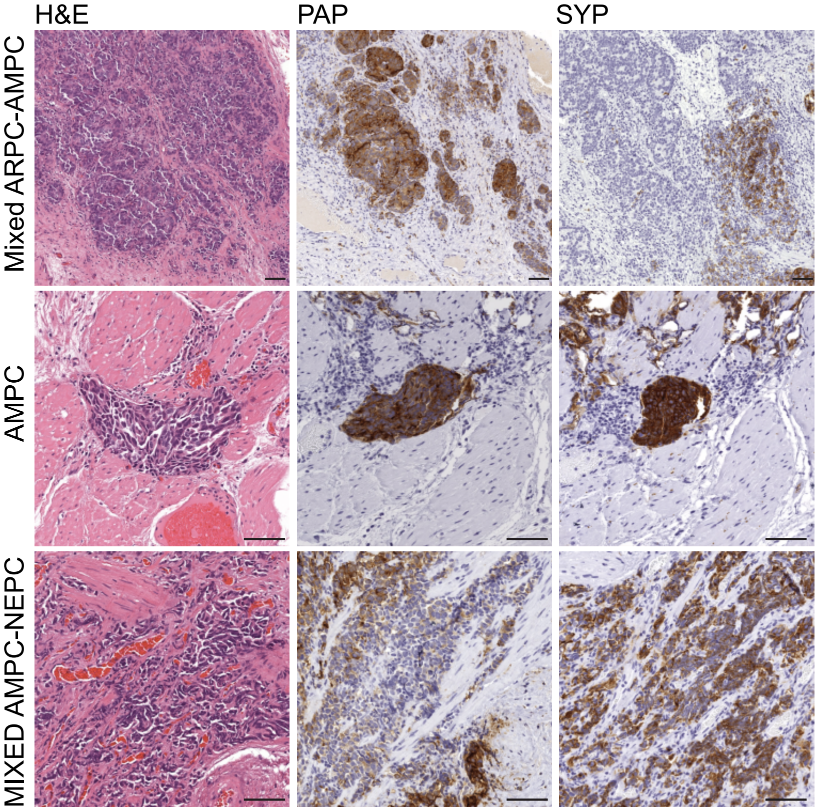


**Supplementary Fig. S11**

In situ profiling of a biopsy specimen of case FH0334 revealed intermixed phenotypes with distinct immunoprofiles and histomorphologies. While the majority of the lesion was composed of cells co-expressing the neuroendocrine marker synaptophysin (SYP) and the AR signaling marker prostatic acid phosphatase (PAP), consistent with amphicrine carcinoma (AMPC), a separate cell population showed only PAP expression, consistent with ARPC. A third cell population demonstrated SYP expression in the absence PAP expression and morphological features consistent with NEPC/small cell neuroendocrine carcinoma. Scale bar indicates 50 um.

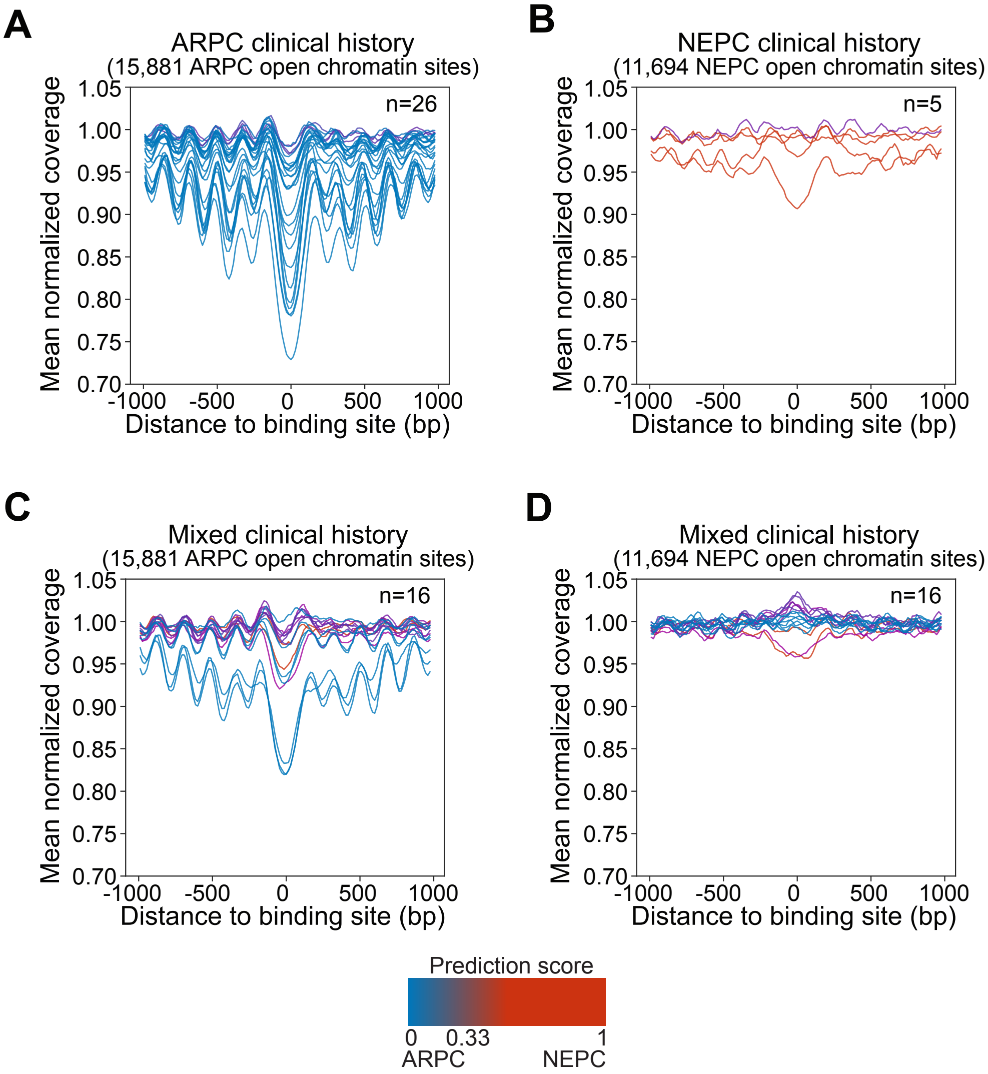


**Supplementary Fig. S12**

**(A)** Griffin cfDNA coverage profiles at ARPC-specific open chromatin sites for the UW cohort samples with ARPC clinical phenotype (n=26). Red, stronger NEPC prediction score; blue, stronger ARPC prediction score; brown, prediction score cutoff threshold of 0.33.

**(B)** (top right) Griffin cfDNA coverage profiles at NEPC-specific open chromatin sites for the UW cohort samples with NEPC clinical phenotype (n=5).

**(C)** Griffin cfDNA coverage profiles at ARPC-specific open chromatin sites for the UW cohort samples with mixed clinical phenotype (n=16).

**(D)** Griffin cfDNA coverage profiles at NEPC-specific open chromatin sites for the UW cohort samples with mixed clinical phenotype (n=16).

**Supplementary Table S1. PDX sequencing metrics**

Phenotype and sequencing metrics for all PDX LuCaP lines.

**Supplementary Table S2. PTM peak data and phenotype 47 fragment variability**

Sheet 1: PDX sample representation in 3 histone PTM CUT&RUN nucleosome profiling assays (H3K4me1, H2K27ac and H3K27me3).

Sheet 2: $Log_2$ fold-change, p-value, and q-value between ARPC and NEPC lines for coefficient of variation in the 47 phenotype defining gene bodies (two tailed Mann-Whitney U test, Benjamini-Hochberg adjusted).

Sheet 3: $Log_2$ fold-change, p-value, and q-value between ARPC and NEPC lines for coefficient of variation in the 47 phenotype defining gene promoters (two tailed Mann-Whitney U test, Benjamini-Hochberg adjusted).

**Supplementary Table S3. Differential regulation metrics and values for select features**

Sheet 1: $Log_2$ fold-change, p-value, and q-value between ARPC and NEPC lines for NPS in the 47 phenotype defining gene bodies (two tailed Mann-Whitney U test, Benjamini-Hochberg adjusted).

Sheet 2: Differentially expressed list of 514 transcription factor (TF). All statistical comparison and fold change estimation was done against ARPCs.

Sheet 3: $Log_2$ fold-change, p-value, and q-value between ARPC and NEPC lines for central mean coverage in 107 TFs overlapping RNA-Seq up/down regulated TFs (two tailed Mann-Whitney U test, Benjamini-Hochberg adjusted).

Sheet 4: Mean values in ARPC, NEPC, and HD lines for central and window means for ARPC and NEPC specific open chromatin regions.

**Supplementary Table S4. Feature-region combination AUCs and benchmarking**

Sheet 1: $Log_2$ fold-change, p-value, and q-value between ARPC and NEPC lines for central mean coverage in all queried (338) TFs (two tailed Mann-Whitney U test, Benjamini-Hochberg adjusted).

Sheet 2: 100-fold cross-validation AUCs for all region and feature combinations subset by the 'AR10' overlapping features (see methods).

Sheet 3: 100-fold cross-validation AUCs for all region and feature combinations subset by the 'Phenotype 47-defining' overlapping features.

Sheet 4: 100-fold cross-validation AUCs for all region and feature combinations (global).

Sheet 5: Predictions scores, tumor fraction, depth of coverage, and subtype for benchmarking admixtures.

Sheet 6: AUCs for unsupervised prediction of admixture subtypes grouped by tumor fraction and depth.

**Supplementary Table S5. Unsupervised model predictions and patient/validation cohort sequencing metrics**

Sheet 1: Histology, tumor fraction, subtype score, and subtype calls for DFCI cohort I.

Sheet 2: Histology, tumor fraction, subtype score, and subtype calls for DFCI cohort II.

Sheet 3: Histology, tumor fraction, subtype score, and subtype calls for WGS and ULP patient samples (UW cohort).

Sheet 4: Complete sequencing metrics for DFCI cohort I.

Sheet 5: Complete sequencing metrics for DFCI cohort II (ULP).

Sheet 6: Complete sequencing metrics for DFCI cohort II (deep WGS)

Sheet 7: Complete sequencing metrics and ichorCNA estimated tumor fractions for the UW cohort (ULP).

Sheet 8: Complete sequencing metrics for the UW cohort (deep WGS)

Sheet 9: Complete sequencing metrics for the healthy donor cohort.
